# Integrative LC-MS and GC-MS Metabolic Profiling Unveils Dynamic Changes during Barley Malting

**DOI:** 10.1101/2024.07.12.603295

**Authors:** Heena Rani, Sarah J Whitcomb

## Abstract

Malting, a crucial process for beer production, involves complex biochemical transformations affecting sensory attributes and product quality. Despite extensive research on storage carbohydrates and proteins involved in malting, a detailed understanding of metabolic alterations during this process remains elusive, limiting our ability to assess and enhance malt quality. Our study employed untargeted GC-MS and LC-MS metabolite profiling to elucidate these changes across six malting stages: dry seed, post-steeping (DOG0), germination (DOG1, DOG3, DOG5), and kilning. We identified a total of 4980 known metabolites, with approximately 82% exhibiting significant changes. Statistical analysis revealed stage-dependent metabolic shifts, with most significant shifts occurring from DOG1 to DOG3 and during kilning. Dynamic changes in various chemical classes and metabolic pathways provide insights into processes critical for malt quality and beer production. Additionally, metabolites associated with antimicrobial properties and stress responses were identified, underscoring the interplay between barley and microbial metabolic processes during malting.

**Highlights:**

- GC-MS and LC-MS profiling were performed to track metabolic changes during malting.
- Identified 4980 known compounds belonging to 346 chemical classes during malting.
- Many microbial metabolites demonstrated increased abundance in finished malt.
- The most significant metabolic shifts occurred during early germination and kilning.

## 1. Introduction

Beer is among the oldest and most widely consumed alcoholic beverages worldwide. Beer production requires the transformation of raw cereal grain into malt, a ready, natural source of nourishment for the yeast used for alcohol generation. Globally, annual malt production ranges from 18 to 22 million tons, 94% of which is used for beer production (Ullrich, 2011). While various grains, including wheat, rye, maize, rice, and oats can be malted, barley (*Hordeum vulgare* L. subsp. vulgare) remains the grain of choice, due to its robust enzymatic activity, beneficial husk properties, and adaptability to diverse climatic conditions (Hamany Djande et al., 2022).

The standard malting process comprises three steps: steeping, germination, and kilning. During steeping, barley grains are soaked in water to raise the moisture content from 12% to 46%. This hydration breaks dormancy and kickstarts the embryo’s early growth, triggers the activation and synthesis of diastatic enzymes (Guido & Ferreira, 2023), and cleans the grain by removing dust/dirt and reducing harmful microbes. Steeping typically lasts 24-48 h, alternating between wet and air rests to meet the increasing demand for oxygen during growth initiation. Once hydrated, excess water is drained, and the barley undergoes germination for 4-5 days in an aerated, humid environment with rotation, resulting in “green malt.” The barley seeds are most metabolically active during the germination stage of malting, with intense enzymatic activity hydrolyzing cell walls and starch, making stored energy accessible for the germinating grain and to yeast during fermentation (Kerr et al., 2023; O’Lone et al., 2023). Following germination, the green malt is kilned with dry air of increasing temperature to reduce grain moisture to 4-5%, halting further germination and ensuring optimal conditions for the conservation and storage of the grain (Daneri-Castro et al., 2016; Rani & Bhardwaj, 2021). Kilning typically lasts 1-2 days, and the maximum air temperature used depends on the malt type being produced: around 80°C for pilsner, 140°C for caramel, and over 200°C for black malts to achieve the desired Maillard reaction products (Iimure & Sato, 2013). The resulting kilned product is the finished malt whose enzymes and metabolites directly contribute to yeast growth during fermentation as well as influence flavor, aroma, and color of beer.

As the barley transforms during malting from a quiescent seed to germinating grain to kilned and dehydrated malt, a myriad of molecular changes occurs in the seed. Shotgun proteomics has detailed the dynamic protein composition throughout malting (Mahalingam, 2018), and several transcriptomics studies have revealed the breadth of gene expression changes across malting stages(Lapitan et al., 2009; Vinje et al., 2021; White et al., 2006). Metabolomics captures the integrated influences of the genome, transcriptome, proteome, and environmental factors, thereby defining the biochemical phenotype of the organism (Fiehn, 2002). However, a comprehensive picture of the barley metabolome across malting stages has remained elusive.

The primary challenge in characterizing the metabolome of malting is the vast chemical diversity of the compounds to be measured and the lack of a single analytical platform to capture that chemical diversity. Prior investigations have tended to focus on a limited array of compounds, such as specific metabolites or subsets of metabolites (Bravi et al., 2012; Beccari et al., 2018; Carvalho et al., 2015; Carvalho & Guido, 2022; Rani et al., 2024; Uygun et al., 2007). Those studies that have sought to more broadly describe the malting metabolome have relied upon a single analytical technique, either nuclear magnetic resonance (NMR) spectroscopy (Byeon et al., 2022), gas chromatography-mass spectrometry (GC-MS) (Frank et al., 2011; Gorzolka et al., 2012) or liquid chromatography-mass spectrometry (LC-MS) (Zhao et al., 2022). Each of these techniques can detect and measure the relative abundance of hundreds to thousands of compounds, but no single technique can come close to capturing the full spectrum of metabolomic changes through the process. Furthermore, some previous metabolomics studies on malting barley have focused on single stages in the malting process (Bettenhausen et al., 2018; Gorzolka et al., 2012; Huang et al., 2016) rather than time-course analysis from dry seed through steeping and germination to kilned product.

This study integrates untargeted GC-MS and LC-MS metabolite profiling and aims to describe the metabolic transformations occurring throughout the malting process. ‘Conrad’ spring malting barley was chosen because this cultivar has been consistently recommended for the past decade to growers and malt producers by the American Malting Barley Association (AMBA) and is used as a check variety for malt quality analysis at the USDA-ARS Cereal Crops Research Unit. The integrative approach employed here bridges critical gaps in the metabolic changes during the malting process and reveals novel insights into the intricate biochemical classes and pathways that are pivotal for determining malt quality and enhancing sensory attributes.

## 2 Materials and methods

For more in-depth details, refer to Supplementary Methods.

### 2.1 Plant materials, malting, and malt quality

Equal quantities of Conrad seed grown in 2014 and 2015 at four barley producing regions were pooled and then malted (Table S1) at the Malt Quality Laboratory of the USDA-ARS Cereal Crops Research Unit (CCRU) in Madison, WI. Briefly, three micro-malting replicates of 110 g each were steeped according to the following regimen: alternating cycles of 4h water immersion at 16°C followed by 4 h air rest at 18°C repeated until the moisture content reached 45%. Post-steeping, seeds were transferred to containers with small sidewall holes and germinated for 120 h at 17°C and > 98% humidity. To prevent matting and to allow gas exchange, the containers were rotated every 30 min for 3 min and moisture levels were maintained at 45%. The germinated seeds were then kilned using the following program: 49°C for 10 h, 54°C for 4 h, 60°C for 3 h, 68°C for 2 h, and 85°C for 3 h with 30 min temperature ramps (Fig. 1). After kilning the moisture content of the finished malt was 4%, the dried culms are removed, and the samples were left to rest for a week to equilibrate the moisture throughout the grains. Malt quality analysis was performed at CCRU using the CCRU methods of analysis of barley and malt (https://www.ars.usda.gov/ARSUserFiles/50900500/barleyreports/CY%20METHODS.pdf) (Table S1).

**Fig. 1:**
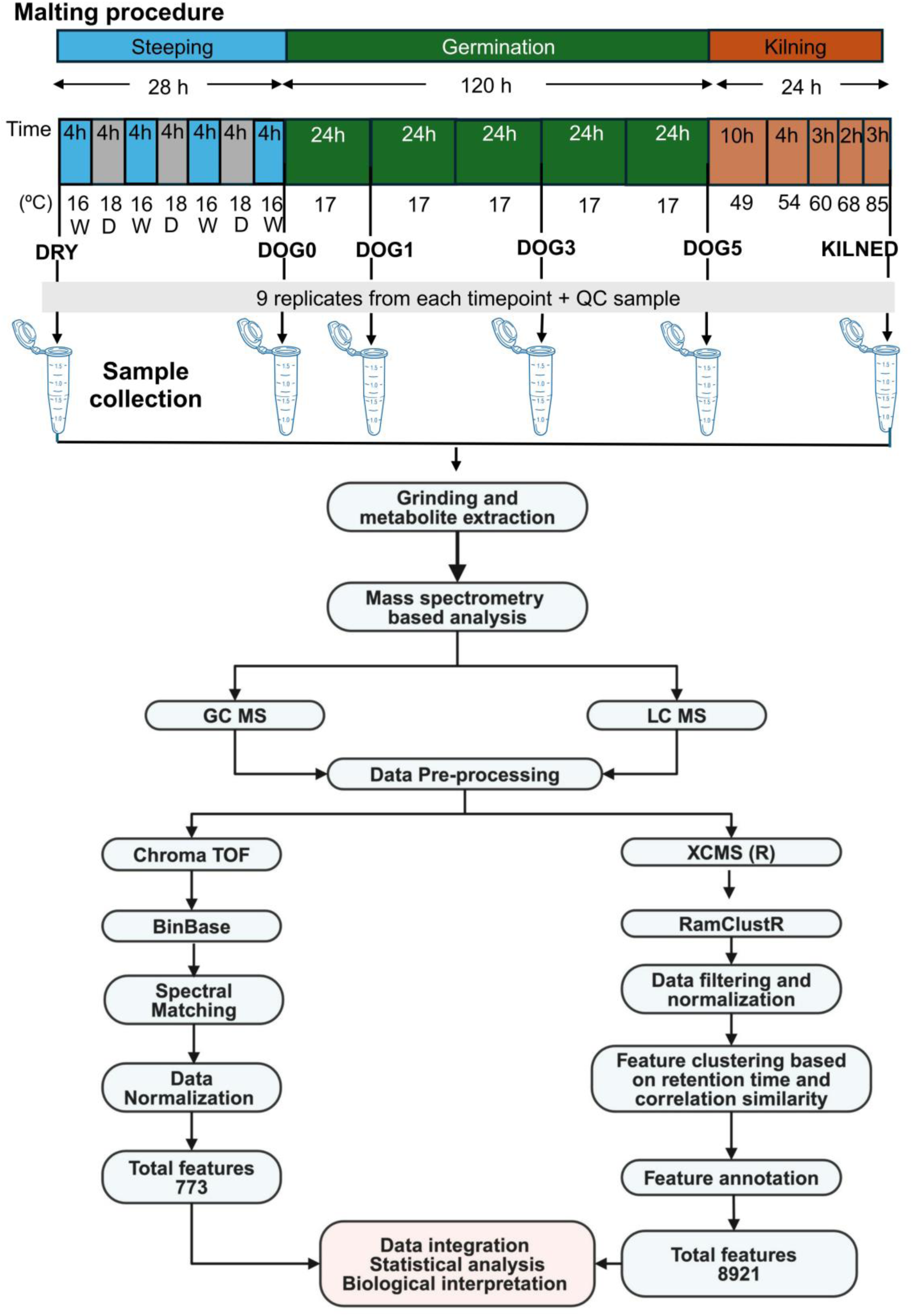
Overview of the malting regime, sample collection, and metabolomics workflow. “W” and “D” indicate wet and air-dry stages of steeping, respectively.

### 2.2 Sample collection and grinding procedure

Nine samples were collected (three per micro-malting replicate) at six different stages: unmalted mature grain (DRY), out of steep/post-step (DOG0), on alternating days of germination (DOG1, DOG3, and DOG5), and at the end of kilning (KILNED) (Fig. 1). Here, “days of germination” refers to the time period after transferring the steeped seeds into rotating chambers for a phase the malting industry calls germination. It is important to note that physiological *sensu stricto* germination is completed by the end of the steeping phase when tiny radicles become visible (referred to as chitting). Therefore, our use of “days of germination” should not be interpreted as the precise time interval post-physiological germination, which occurs during the steeping stage.

Samples were collected in Eppendorf tubes, immediately flash frozen in liquid nitrogen and subsequently stored at −80°C until grinding. For grinding, the seeds were lyophilized, placed in 5 ml polycarbonate vials along with three stainless steel balls (6.3 mm), and cooled in liquid nitrogen. The frozen tissue was ground to a fine powder in a 2010 Geno/Grinder (Spex SamplePrep) with intermittent rests, rapidly re-cooled in liquid nitrogen, and transferred to their final storage vials. The ground samples were stored at −80°C until further analysis.

### 2.3 Metabolite extraction

#### 2.3.1 Preparation of samples for GC-MS analysis

Finely ground samples (4 mg) were extracted using 1 ml of 3:3:2 isopropanol (IPA)/acetonitrile (ACN)/water (v/v/v) for 6 min at 4°C. After centrifugation, the supernatant was aliquoted into two equal portions and dried. One aliquot was derivatized with methoxamine, a mixture of fatty acid methyl esters (FAMEs) ranging from C08 to C30 was added to each sample, and finally derivatized with N-methyl-N-trimethylsilyltrifluoroacetamide (MSTFA). The prepared samples were then transferred to vials, sealed, and loaded into the GC-MS instrument. Quality control (QC) samples were prepared by pooling equal amounts of ground powder from all biological samples and were processed as described above. Method blanks were prepared using the same 3:3:2 IPA/ACN/water mixture without the addition of biological sample and processed using the same extraction and processing procedures as the other samples.

#### 2.3.2 Preparation of samples for LC-MS analysis

Finely ground samples (100-130mg) were washed with 800 μl of 80% IPA in water for 10 min at room temperature. After centrifugation, 700 μl of the supernatant was removed and discarded. This process was repeated with 700 µl of 80% IPA. The pellet was then extracted twice using 700 µl of 80% methanol. The 1.4 ml of combined methanol extract was dried under nitrogen and then reconstituted in 100 µl of methanol. The reconstituted extracts were then transferred to autosampler vials for UPLC-MS analysis. QCs were prepared by equally pooling ground powder from all samples and were processed as described above. Method blanks were prepared and processed in the same manner without the addition of biological sample.

### 2.4 Metabolomic Fingerprinting

#### 2.4.1 GC-MS measurements

GC-MS measurements were performed on an Agilent 7890A GC coupled with a Leco Pegasus IV time-of-flight mass spectrometer (LECO Corporation, St. Joseph, MI, USA). Derivatized samples were injected using a splitless method into a multi-baffled glass liner. Pooled QCs were injection after every 10 samples. Compounds were separated using the GC-Column DB-5MS with an Intergra-Guard and a helium flow rate of 1 mL/min. The electron ionization (EI) energy was set to 70 eV. The system collected the mass spectra in a mass range of 85-500 Da at an acquisition rate of 17 spectra/s.

#### 2.4.2 LC-MS measurements

LC-MS measurements were performed on a Waters Acquity UPLC system (Waters Corporation, Milford, MA, USA) with a pooled QC injection after every five samples. Compounds were separated using a Waters Acquity UPLC CSH Phenyl Hexyl column (1.7 μM, 1.0 x 100 mm) with a gradient from 0.1% ammonium formate in water to 0.1% formic acid in acetonitrile with a 200 μl/min flow rate. The column eluent was infused into a Waters Xevo G2-XS Q-TOF-MS with an electrospray source in positive mode, scanning 50-1200 m/z at 0.1 seconds per scan, alternating between MS (6 V collision energy) and MSE mode (15-30 V ramp).

### 2.5 Data treatment and metabolite identification

#### 2.5.1 GC-MS

Following data collection, raw data files were preprocessed using Leco ChromaTOF software v2.32 with baseline subtraction just above the noise level and automatic mass spectral deconvolution and peak detection at a signal/noise ratio of 5:1. Apex masses were reported for use in BinBase algorithm (Fiehn et al., 2005). Result files were exported to a data server with absolute spectra intensities and further processed by a filtering algorithm implemented in the metabolomics BinBase database. Spectra were cut to 5% base peak abundance and matched to database entries from most to least abundant spectra using the following matching filters: retention index window ±2,000 units (equivalent to about ± 2 s retention time), validation of unique ions and apex masses (unique ion must be included in apexing masses and present at > 3% of base peak abundance), mass spectrum similarity must fit criteria dependent on peak purity and signal/noise ratios and a final isomer filter. Raw data peak heights were normalized to the sum of peak heights of all identified metabolites in each sample. One of the replicates from the dry seeds failed and was removed from further GC-MS analysis.

#### 2.5.2 LC-MS

R package XCMS v3.20.0 (Smith et al., 2006; Tautenhahn et al., 2008)was used to process raw data. XCMS steps included: peak detection (CentWave), peak grouping (PeakDensity), retention time correction (PeakGroups), peak regrouping (PeakDensity), and missing peak filling (FillChromPeaks).

R package RAMClustR v1.2.4 (Broeckling et al., 2014)was used to normalize, filter, cluster features into spectra, and infer molecular weights. Missing values were replaced with small values simulating noise: for each feature, the replacement value was equal to the absolute value of 0.5 times the minimum detected value of that feature. Features were normalized by linearly regressing run order versus QC feature intensities to account for instrument signal intensity drift. Only features showing significant regression (p < 0.05, r-squared > 0.1) were corrected. Features failing to demonstrate at least a 2-fold greater signal intensity in QC samples compared to blanks were removed. Subsequent filtering based on QC sample coefficient of variation (CV) values retained only features with CV values ≤ 0.5 in MS or MSMS data sets. Features were clustered using the ramclustR algorithm (Broeckling et al., 2014).

MSFinder v 3.52 (Tsugawa et al., 2016)was used for spectral matching, formula inference, and tentative structure assignment. MSFinder results were imported into the RAMClustR object where a total score was computed based on the product scores from the findmain function and the MSFinder formula and structure scores. Spectra matches took precedence over computational inference-based annotations. CHEBI and COCONUT were set as priority databases, and matches to these databases were given a priority factor value of 1. Matches to databases other than the priority databases were assigned a priority factor of 0.9. Based on associations from FooDB and PubChem, a custom database of barley compounds was generated. The list of 8858 InChIKey structures in this custom barley database was also used for annotation prioritization. Annotations with InChIKey(s) that did not match those in the custom barley database were assigned an InChIKey priority factor of 0.9 to decrease scores for these annotations. The annotation with the highest total score was selected for each compound.

### 2.6 Dataset Integration and Statistical Analysis

We mainly focused on known metabolites (those with InChIKeys). Nearly 7,600 out of 8,921 LC-MS features and 170 out of 773 GC-MS features corresponded to known metabolites (Table S2-3). To include features with high confidence, we excluded the LC-MS features from different clusters with identical annotations. In cases, where a metabolite was detected in both datasets, we selected the data from the platform with the lower pool QC CV value (Table S4). Final integrated dataset comprised 4,980 known metabolites (Table S5).

Data normalization was conducted using Box–Cox transformation (Yu et al., 2022)and assessed using the Shapiro-Wilk test. For metabolites with normal distribution, Levene’s test was used to assess the homogeneity of variances. Metabolites with unequal variances (p < 0.1) were analyzed using Welch’s t-test and Welch’s ANOVA, while those with equal variances were subjected to Student’s t-test and standard ANOVA. Metabolites not following a normal distribution after Box-Cox transformation (p < 0.1) underwent Mann–Whitney U test for comparison between consecutive stages and the Kruskal-Wallis test for ANOVA. Significantly differential metabolites met the following two criteria: (i) false discovery rate (FDR) < 0.05, and (ii) fold change ≥ 2 or ≤ 0.5.

Compounds were assigned to chemical ontology using ClassyFire (Djoumbou Feunang et al., 2016). Chemical similarity enrichment analysis (ChemRICH) was performed using raw p-values derived from the different statistical tests applied for comparing means in consecutive stages (such as the Mann-Whitney U test, Welch’s t-test, and Student’s t-test) and parent class level 1 information as set definition. Sets having a frequency of less than three and sets having no compounds with a raw p-value < 0.10 were excluded. Sets with FDR < 0.01 were considered significant. PCs were grouped based on their respective superclass, and hierarchical clustering analysis (HCA) was performed on the -log10(FDR) values obtained from the ChemRICH analysis. Annotated metabolite names were converted to KEGG IDs using various platforms: Refmet (Fahy & Subramaniam, 2020), Mbrole2.0 (López-Ibáñez et al., 2016), Chemical Translation Service (Wohlgemuth et al., 2010), PubChem Exchange Identifier service (https://pubchem.ncbi.nlm.nih.gov/idexchange/), and Metaboanalyst 6.0 (Pang et al., 2024). Out of 4,980 annotated metabolites in the dataset, 600 corresponded to unique metabolite-KEGG ID pairs. The data was auto scaled and Kyoto Encyclopedia of Genes and Genomes (KEGG) pathway analysis was performed. MetaboAnalyst 6.0 (Pang et al., 2024)was used to identify and visualize the altered metabolic pathways using *Oryza sativa* japonica as the background set. The global test and relative-betweenness centrality were used for the enrichment method and pathway topology analysis, respectively. The significantly affected pathways (with at least five metabolites detected in our dataset) were selected by FDR < 0.05 from pathway enrichment analysis.

### 2.7 Raw data

This study is available at the NIH Common Fund’s National Metabolomics Data Repository (NMDR) website, the Metabolomics Workbench(Sud et al., 2016), https://www.metabolomicsworkbench.org where it has been assigned Study ID ST003289. The data can be accessed directly via its Project DOI: https://dx.doi.org/10.21228/M8SN77. The NMDR is supported by NIH grant U2C-DK119886 and OT2-OD030544 grants.

## 3 Results and discussion

### 3.1 Untargeted metabolite profiling of six malting stages using LC-QTOF-MS and GC-TOF-MS

Our LC-MS approach detected 8,921 features (7,600 knowns) (Table S2) in the malting time course samples, and GC-MS detected 773 features (172 knowns) (Table S3), with 38 known compounds (Table S4) being common to both analytical platforms (Fig. 1, 2A). Intrinsic differences exist in separation and detection capabilities of LC-MS and GC-MS platforms, each with its unique advantages. GC-MS is more suitable for volatile, thermally stable, and small polar metabolites covering primary metabolism, whereas LC-MS can analyze non-volatile, thermolabile, and more complex hydrophobic metabolites predominant in secondary metabolites (Doerfler et al., 2013).

**Fig. 2:**
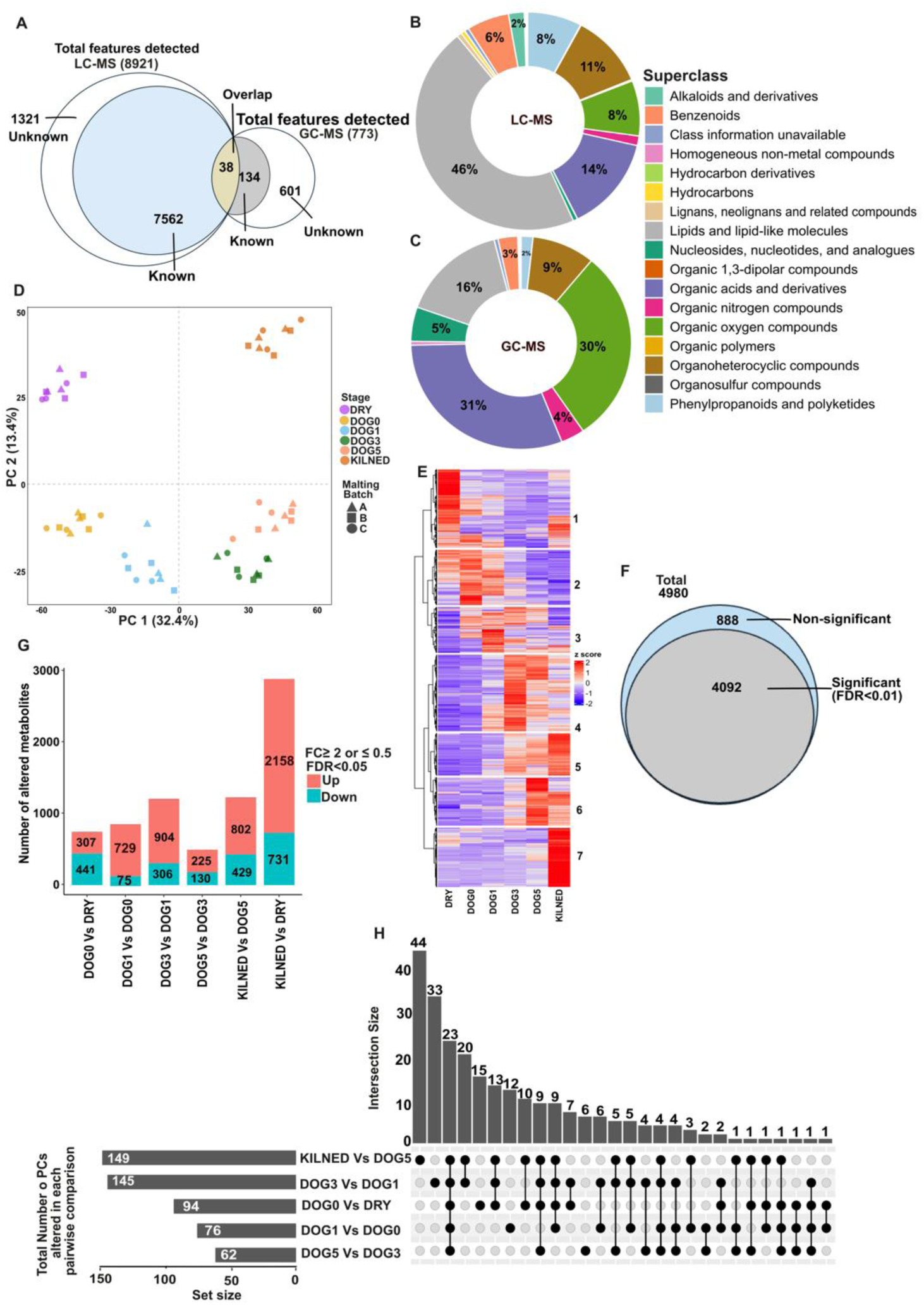
(A) Number of features detected by LC-MS and GC-MS, the subset of those that were annotated to known metabolites, and the unknown metabolites detected by both analytical platforms. (B, C) Classification of known metabolites in the LC-MS and GC-MS datasets based on ClassyFire superclass. (D - H) Analysis on the integrated LC-MS and GC-MS dataset comprising 4980 metabolites. (D) Scores plot of principal component analysis, performed after Box-Cox transformation and autoscaling. Different symbols indicate the three distinct malting batches, each with three replicates, while colors represent the malting stage. (E) Hierarchical clustering analysis using the Euclidean distance metric and the Ward.D2 method, applied to mean centered and standardized abundance data. (F) Number of significantly altered and non-significantly altered metabolites during malting based upon ANOVA (FDR < 0.01). (G) Number of significantly altered metabolites between the indicated stages based on different statistical tests applied for comparing means in consecutive stages (as mentioned in 2.6). (H) Set analysis on parent classes significantly altered at each stage (FDR < 0.01).

Metabolites with known InChIKeys were categorized based on chemical structural features via ClassyFire (Djoumbou Feunang et al., 2016). The distribution of metabolites across different chemical structural class levels is summarized in the interactive pie chart shown in Fig. S1. The LC-MS dataset comprised compounds from 15 super classes (Fig. 2B), and the GC-MS dataset contained compounds from nine (Fig. 2C). In the LC-MS dataset, 46% of the known metabolites are lipid and lipid-like molecules, with prenol lipids being the dominant class. The remaining 54% belong to diverse chemical classes, including 1,076 organic acids and their derivatives (OADs), 832 organoheterocyclic compounds (OHCs), 627 organic oxygen compounds (OOCs), 596 phenylpropanoids and polyketides (PPs), 467 benzenoids, and 172 alkaloids and derivatives (Fig. 2B). GC-MS was less effective in detecting large, non-volatile molecules like lipid and lipid-like molecules (16%), but more suited for smaller, volatile compounds like organic oxygen compounds, organic acids, and organic acid derivatives, comprising 61% of the known metabolites with InChiKeys. The remaining 23% include 16 OHCs, nine nucleosides, nucleotides and analogues, and six organic nitrogen compounds (Fig. 2C).

PCA was conducted on the GC-MS and LC-MS datasets separately. The QC samples in both datasets grouped tightly, and replicates from each malting stage clustered distinctly, indicating low intra-group variation relative to inter-group variation and overall robustness of the instruments and post-acquisition data processing (Fig. S2). After filtering out unknowns and duplicate features from the initial dataset of 8,921 features (LC-MS) and 773 features (GC-MS), the integrative approach resulted in a final dataset of 4,980 known metabolites (Table S5). This combined dataset was also subjected to PCA (Fig. 2D). PC1 accounted for 32.4% of the total variance and separated the end of steep (DOG0) to end of germination (DOG5) sample groups in a unidirectional manner. The position of samples from the six malting stages on the PCA scores plot reflect the metabolic journey, starting from a dormant state, moving through a phase of high metabolic activity and diversity during germination, and then becoming less metabolically active during kilning. This U-shaped pattern has also been observed in previously published metabolomics studies of barley malting using LC-MS (Zhao et al., 2022) and GC-MS (Frank et al., 2011).

Hierarchical Clustering Analysis (Fig. 2E) classified metabolites into 7 clusters based on their z-score across malting stages. Metabolites in clusters 1, 2, and 3 (44%) exhibited higher abundance in the initial phase of malting (up to DOG1). Compounds including lignin, lignans, neolignans and related compounds, as well as various phenylpropanoids and polyketides predominate in these early accumulating clusters. The increase in these metabolites during early stages of malting is likely due to the enzymatic breakdown of cell wall constituents and the release of phenolics bound to cellular structures (e.g., lignin and arabinoxylans) (Quan et al., 2018). The phenylpropanoid and polyketides included metabolites with antimicrobial properties, as discussed in detail in sections 3.2.2 (b) and (c).

Clusters 4, 5, 6, and 7 (56%) comprised metabolites that peaked between DOG1 and Kilning. Clusters 5 and 7 contained various heterocyclic compounds whose abundance increased dramatically during kilning. These compounds contribute to organoleptic character of beer and are likely formed by degradation of higher-structure molecules or through non-enzymatic interactions between free amino acids and reducing sugars during kilning (Prumysl et al., 2021; Svoboda et al., 2022).

### 3.2 Differential Abundance Analysis

#### 3.2.1 ANOVA and Pairwise comparisons

One-way ANOVA revealed significant changes in abundance for about 82% of the metabolites in our dataset during malting (ANOVA FDR > 0.01) (Fig. 2F). These metabolites may be regulated homeostatically or they may not be directly involved in malting transformations. To track metabolic changes among the six malting stages, statistical tests (as mentioned in 2.6) were performed and fold changes were calculated on data from consecutive stages and between dry (malting input) and kilned (malting output) samples, resulting in six comparisons (Fig. 2G). There was an increase in 2,158 metabolites and a decrease in 731 in finished malt compared to dry seed. The transitions from DOG1 to DOG3 and from DOG5 to KILNED had the largest number of differentially abundant metabolites. Between DOG1 to DOG3, many hydrolytic enzymes that contribute to the modification of the physical and chemical structures of the kernel are expressed (Betts et al., 2017; Leišová-Svobodová et al., 2020; Schmitt et al., 2013; Vinje et al., 2021). These enzymes facilitate the degradation of cell walls, depolymerization of starch, and breakdown of lipids and proteins during the early stages of germination, resulting in a vast number of metabolic changes. (Table S5). During kilning, many metabolic changes are likely driven by high temperatures that promote the Maillard reaction, which is responsible for an increased abundance of various Maillard reaction products, contributing to the complexity of malt flavor (Briggs, 1998). The decrease in various metabolites during this stage is likely due to heat-induced degradation or their utilization in the Maillard reaction. Of the five comparisons between consecutive malting stages, the transition from DOG3 to DOG5 showed the fewest metabolites with significantly altered abundance (255 up, 130 down), suggesting a possible stabilization in metabolic activity. This apparent stabilization is likely a result of inhibited growth due to several factors including the rotation (anti-gravity effects), crowding (thousands of seedlings in close proximity), and high humidity.

In all the studied comparisons, the number of metabolites that increased in abundance exceeded the number that decreased, except during steeping (Fig. 2G). The larger number of decreasing metabolites during steeping is likely due to their leaching from the barley grains into the steep-out water, including substances from the outer layers of the grain (pericarp and testa), metabolites from microbial sources, and those loosely associated with cellular structures. Meanwhile, the compounds that increased during steeping, such as various hydrolytic products, amino acids, and antioxidants, likely result from the activation of biochemical processes triggered by water absorption.

#### 3.2.2 Chemical Enrichment Analysis

To assess alterations in chemical classes during malting, ChemRICH analysis was conducted. This method identifies highly impacted compound classes by clustering metabolites based on chemical similarity and ontologies, enabling identification of biologically relevant groups of metabolites that may not be annotated in traditional pathway databases(Barupal & Fiehn, 2017). Additionally, it does not rely on background databases for statistical calculations. Parent Classes (PCs) with at least three compounds in our dataset or with at least one compound having a raw p-value < 0.1 in the relevant differential abundance test between consecutive malting stages were tested by ChemRICH. 4470 compounds classified into 346 PCs met these criteria, with 243 PCs showing significant alterations (FDR < 0.01) in at least one comparison between malting stages (Table S6, S7).

Of the 243 significantly altered PCs, 94 were significantly altered during steeping, with most showing decreased abundance of metabolites likely due to leaching. Among these 94 PCs,15 are unique to steeping (Table S8, Fig. 2H). During the entire germination phase, 174 classes were significantly altered, with 67 being unique to germination. Among these 67 unique classes, 33 were specifically altered during the transition from DOG1 to DOG3, suggesting distinct metabolic shifts occur during this specific phase. Kilning resulted in significant changes in 149 classes, with 44 unique to this stage (Fig. 2H). Among these, 19 PCs are OHCs (Table S8), known for their flavor-active properties in malt (Coghe et al., 2004; Fors, 1983). These findings align with a recent study showing increased heterocyclic compounds in all (pseudo)cereals after kilning (Almaguer et al., 2023). PCs were grouped based on their respective superclasses, and hierarchical clustering analysis (HCA) was performed on the - log10(FDR) values obtained from the ChemRICH analysis (Table S9). This approach facilitates visualization of the most significantly altered metabolite parent classes within each superclass. The following sections discuss the ChemRICH results for each superclass.

##### 3.2.2.1 OOCs

Among the 4470 metabolites used in ChemRICH analysis, 493 belonged to the superclass OOCs. The 14 PCs belonging to this superclass divided into three clusters (Fig. 3A). Glycosyl compounds and aminosaccharides in cluster 1 showed significant changes at each malting stage. About half of the metabolites in these PCs increased in abundance by the end of malting (Table S9). The increase in simple glycosyl compounds such as maltose, beta-gentiobiose, trehalose, isomaltose, and lactose, as well as complex glycosyl compounds during the initial stages of malting, is likely due to the degradation of cell walls and release of stored reserves by active hydrolytic enzymes (α- and β-amylases, α-glucosidase, and limit dextrinase) (Faltermaier et al., 2015). These hydrolytic processes are critical for successful barley malting, and as such contribute to important malt quality parameters (American Malting Barley Association Inc, 2019). Increase in many glycosyl compounds during kilning is likely due to relatively heat stable glycosyltransferases (Zhao et al., 2022). Interestingly, most of the 116 aminosaccharides detected were antibiotics, indicating the presence of endophytic bacteria and fungi during malting (Table S7). Among the most strongly increasing antibiotics in the kilned malt were Amaromycin, gentamycin C1a, Boholmycin, Fortimicin B, Kanamycin B, as well as many lesser studied compounds. Although microbial load has been observed to decrease by 10 to 100-fold during kilning (Lenaerts et al., 2011), certain heat-resistant fungi continue to grow, possibly forming biofilms that confer thermal resistance (Douglas & Flannigan, 1988; Laitila et al., 2011)

**Fig. 3:**
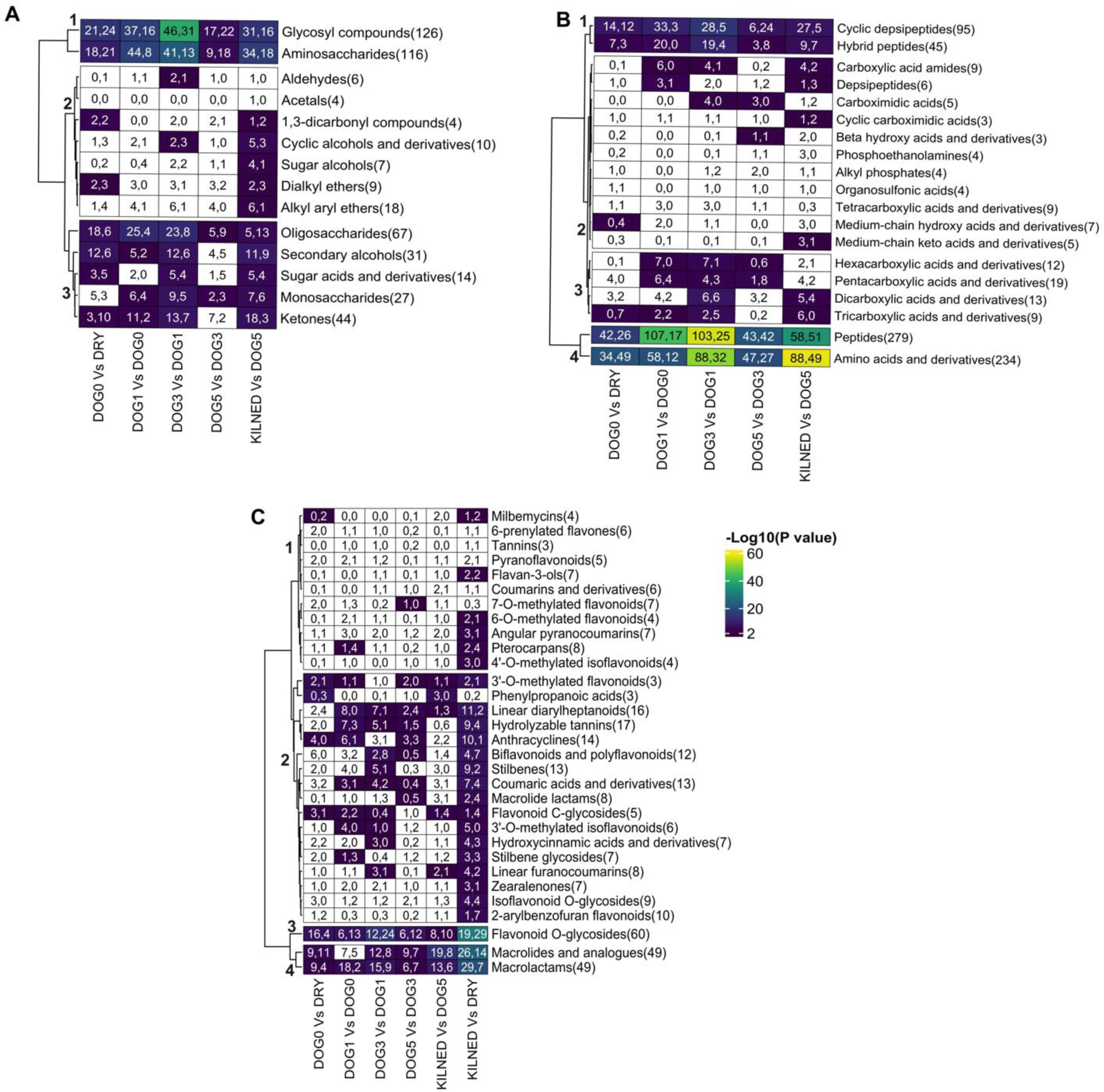
Hierarchical clustering analysis of ClassyFire parent classes based on ChemRICH FDR in the five indicated stage comparisons. The analysis was conducted using the Euclidean distance metric and the Ward.D2 clustering method. Color represents -log_10_(FDR) values. White indicates FDR > 0.01. Each cell contains two numbers, separated by a comma, corresponding to the number of metabolites in that parent class that significantly increased and that significantly decreased, respectively. The total number of metabolites detected in each parent class is indicated in parentheses. (A) Parent classes belonging to organo oxygen compounds superclass, (B) Parent classes belonging to organic acids and derivatives superclass, (C) Parent classes belonging to polyphenols and polyketides superclass.

In cluster 2, PCs such as 1,3-dicarbonyl compounds, cyclic and sugar alcohols, alkyl, aryl, and dialkyl ethers showed significant changes during kilning. Some enzymatic activities might persist early in kilning, until higher temperatures are reached, transforming substrates into alcohols. The decrease in aldose sugars (glucose, xylose, xylulose, lyxose) with concurrent increase in sugar alcohols (sorbitol, xylitol, lyxitol) (Table S7) indicates oxidative thermal decomposition of sugars or enzymatic isomerization during kilning. The high abundance of the enzyme xylose isomerase, which converts xylose to xylulose, was reported during kilning (Mahalingam, 2018).

All cluster 3 PCs displayed significant alterations during the transition to DOG3 and during kilning. Ketones contribute to the aroma and flavor of malt, and the abundance of 26 of the 44 measured ketones increased by the end of malting (Table S9), likely due to thermal degradation of proteins and fatty acids, deamination of amino acids, and decarboxylation of carboxylic acids during kilning. Additionally, the decrease in various secondary alcohols (Table S9) during kilning suggests their oxidation to ketones. The accumulation of many oligosaccharides during steeping and early germination suggests that the breakdown of larger polysaccharides and cell walls begins during steeping and increase during the initial phases of germination (Fig. 3A). For example, maltotriose levels were observed to increase during steeping and DOG1 and remained unchanged at following consecutive stages in the present study. Presence of maltotriose in steep out barley samples and its subsequent increase, beginning from DOG1, was also reported in a prior study (Vinje et al., 2015). Scanning electron microscope images of barley starch in steep out samples and by the end of DOG1 showed no surface changes, while submicron holes appeared in some smaller granules by DOG2 (Contreras-Jiménez et al., 2019). Based on these observations, we speculate that enzyme activities and concomitant changes in metabolite levels may initiate earlier during steeping and DOG1 while their manifestations leading to the ultrastructural changes in starch granules occur in later stages.

##### 3.2.2.2 OADs

Peptides (279) and amino acid derivatives (234) were the most common OAD PCs (clusters 4 and 5) (Fig. 3B), and over half of the associated metabolites increased in the final malt (Table S9). Metabolites in these PCs increase during steeping and showed highly significant rises between DOG0 and DOG3, indicating peak protease activity in early germination, perhaps to supply amino acids for new protein synthesis and to feed the TCA cycle to generate energy (Byeon et al., 2022; Frank et al., 2011; Kim et al., 2020). Furthermore, increases in various amino acids and peptides were also observed during kilning, aligning with the detection of various enzymes with proteinase activities, such as carboxypeptidase, cysteine peptidase and prolyl aminopeptidase in the kilned samples (Mahalingam, 2018). Decreases in the abundance of many peptides and amino acids during kilning are likely due to formation of melanoidins, which are essential for malt color, flavor, and aroma. The results suggest that the extent of protein and amino acid transformation during early germination impacts malt and beer quality.

In addition to peptides and amino acid derivatives, many cyclic depsipeptides (CDP) were detected (cluster 1). CDPs, characterized by a cyclic peptide backbone and ester linkages (Taevernier et al., 2017), are synthesized by both prokaryotic and eukaryotic organisms and exhibit various bioactivities, including antibacterial and antifungal properties (X. Wang et al., 2018). Here, 96 CDPs were identified, with 54 showing increasing abundance and 12 decreasing (FDR < 0.05) in the finished malt. Most of these increased during first three days of germination (cluster 4, Fig. 4A). Although the steeping process reduces surface microbes through repeated cycles of water immersion and air rest, it is unlikely to significantly reduce internal grain microbes. Moisture imbibed during steeping may allow microorganisms to proliferate during germination, leading to increased production of various antimicrobial compounds like CDPs to protect their niche by killing competitors and modulating symbiotic interactions. The abundance of some CDPs also increased during kilning, indicating that the microbial community surviving the initial stage of kilning may continue producing CDPs. While some CDPs act as biocontrol agents, others present challenges to cereal-based industries, including baking, malting, and brewing. For example, enniatins, cyclic hexadepsipeptides produced by Fusarium species, are major contaminants of cereal grains (Gautier et al., 2020; Gnonlonfoun et al., 2023). In this study, five enniatin variants (Enniatin A, A1, B, B1, and K1) were detected, and all showed increased levels in finished malt albeit with relatively high p-values (Fig. 4A). Fusarium growth has been reported during steeping even when the original barley had low-level contamination (Lenaerts et al., 2011). Additionally, various hybrid peptides produced by microbes increased during early germination. The role of CDPs and hybrid peptides in malting processes and their impact on malt quality are not well understood, indicating a crucial need for further detailed studies to explore their effects on malt quality.

**Fig. 4:**
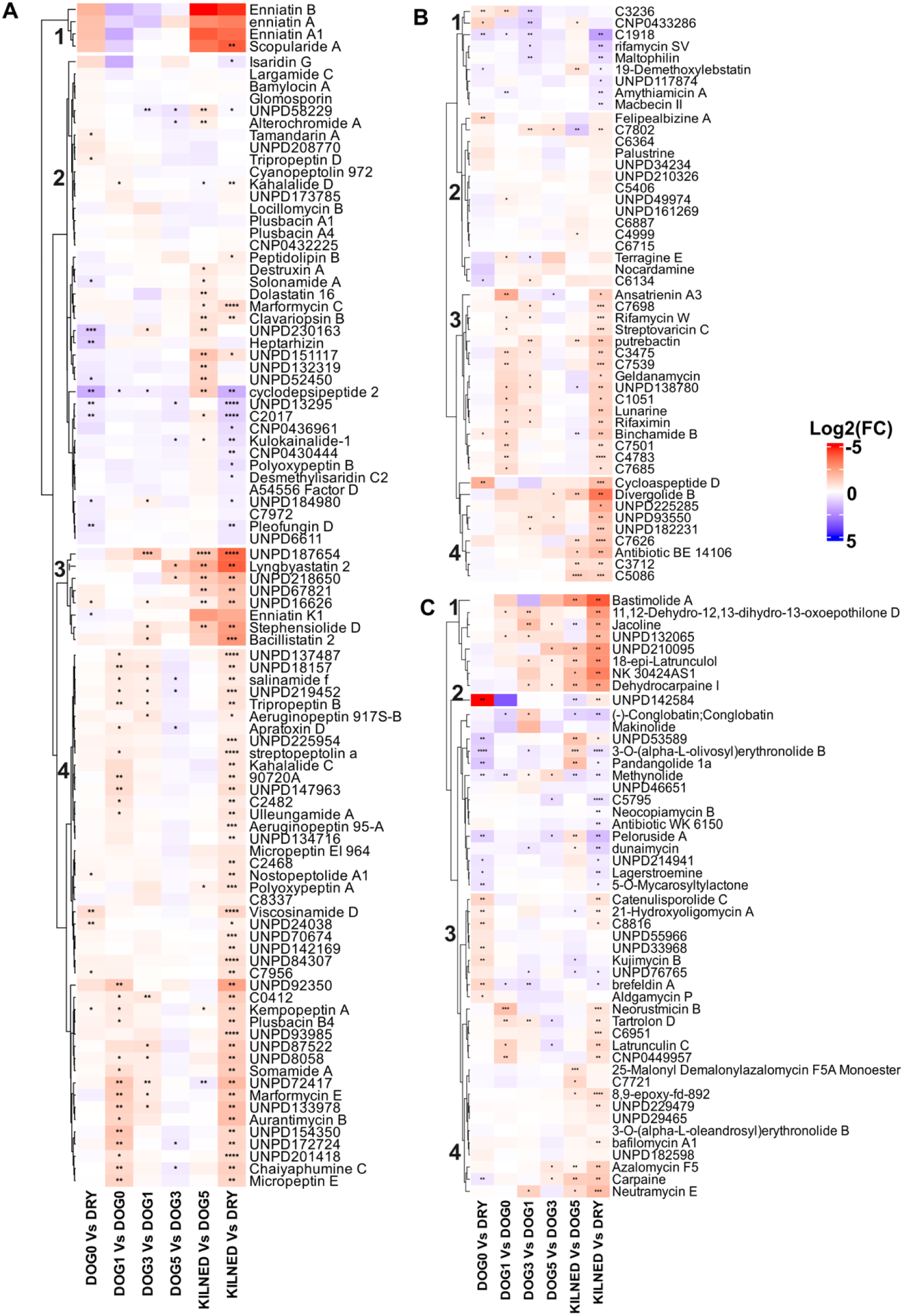
Hierarchical clustering analysis of fold changes between stages in the six indicated comparisons for metabolites in the (A) cyclic depsipeptides parent class, (B) macrolactams parent class, (C) macrolides and analogues parent class. The clustering analysis was conducted using the Euclidean distance metric and the Ward.D2 clustering method. Color indicates the log2(fold change). Statistical significance is indicated by ‘*’ for FDR ≤ 0.05, ‘**’ for FDR ≤ 0.01, ‘***’ for FDR ≤ 0.001, and ‘****’ for FDR ≤ 0.0001. Metabolites with lengthy names are labeled by their cluster ID (Tables S5).

##### 3.2.2.3 PPs

Numerous microbe-derived metabolites belonging to the superclass PP were also identified (Fig. 3C), macrolactams, macrolides and analogues (macrocyclic polyketides). Macrolides, with their large macrocyclic lactone ring, are synthesized by bacteria and fungi, while macrolactams, large cyclic amides with a lactam ring, are predominantly produced by bacteria and rarely found in fungi (Zhou et al., 2022). Our results demonstrate dynamic changes in the concentrations of these microbial classes throughout the barley malting process. Like CDPs, a majority of the macrolactams that increased in final malt showed significant accumulation during first three days of germination (cluster 3, Fig. 4B), whereas macrolides and their analogues showed the most pronounced changes during kilning (Fig. 4C). Most anthracyclines and linear diarylheptanoids also increased in final malt (Table S9). Zearalenones and milbemycins, mycotoxins produced by Fusarium and Streptomyces species, respectively, were also detected and several significantly accumulated in kilned malt relative to dry grain (Table S9). In contrast to our findings, Piacentini et al., (2019) observed a decrease in ZEN levels during the transition from barley to malt and beer. The authors reported that most of this mycotoxin was transferred to the rootlets and spent grains. These differences in results may arise from differences in the barley genotypes used and their growth conditions or from the specific malting regimes employed.

By the end of malting, many compounds from microbial metabolite classes had accumulated. Future research should aim to further characterize these microbial metabolites under various malting conditions to elucidate their roles (including antibiotic production, toxins, enzyme catalysis, biocontrol, and signaling) and optimize their beneficial properties. Understanding these dynamics could lead to improved malting strategies, enhancing both the safety and quality of malt. Additionally, antimicrobial compounds could serve as innovative biomarkers for microbial quality assessment in malting, potentially transforming industry practices and ensuring higher standards of product integrity.

Aside from microbial metabolites, several phenolic classes were identified within the superclass PP (Fig. 3C). There was no general trend observed in phenolic classes; some remained relatively unchanged through malting, while others predominantly increased during the early germination stage. Classes such as conjugated flavonoids (including stilbene and flavonoid O- and C-glycosides), biflavonoids, and polyflavonoids showed prominent reductions during germination. The observed decrease in glycosylated phenols might be due to the activation of β-glucosidase during germination. Conversely, certain polyphenols increased during early germination, likely due to the enzymatic breakdown of cell wall constituents and the release of phenolics bound to cellular structures (Quan et al., 2018). For example, ferulic acid, which accounts for more than 70% of the phenolic acids (Hefni et al., 2019), increased by 2.2-fold from DOG1 to DOG3, potentially due to the liberation of its bound form from cell wall constituents by the enhanced activity of feruloyl esterase during germination. However, catechin levels decreased likely due to its glycosylation during this stage (Table S5) (Carvalho & Guido, 2022). In agreement with the current study, previous studies reported the increase in ferulic acid and the decrease in catechin during germination in barley malt (Friedrich & Galensa, 2002; Leitao et al., 2012). Subsequently, ferulic acid content decreased (1.7-fold) during kilning, perhaps due to the incorporation of phenolic compounds into the structure of melanoidins. This reduction is beneficial as the thermal degradation of ferulic acid can lead to the formation of p-vinylguaiacol through decarboxylation, which may introduce off-flavors to the malt (Naim et al., 1988). Published reports vary regarding total phenolic content changes during malting stages, potentially due to differences in malt origin or malting conditions (Gallegos-Infante et al., 2010; Ha et al., 2016; Pejin et al., 2009). It has been suggested that later stages are marked by lignification and conversion of phenolic compounds to lignans or lignin (Ha et al., 2016). However, our findings did not show a significant increase in lignans, neolignanes, and related compounds at DOG5 (Fig. S3A).

##### 3.2.2.4 OHCs, Benzenoid, Alkaloids and derivatives

Most OHCs increased by the end of malting, especially during kilning (Table S9), likely due to thermal reactions, cyclization, and interactions with reactive molecules like amines and aldehydes (Bettenhausen et al., 2020). These compounds significantly contribute to the flavor and color of malt based on their structure and originating amino acids (Coghe et al., 2004). Here, the most abundant OHCs were nitrogen-based heterocycles, with some containing sulfur and nitrogen (e.g., naphthothiazoles, benzothothiazoles, phenothiazines) or oxygen and nitrogen (e.g., quinolines, quinolones, oxepins, benzodioxoles) (Fig. 5).

**Fig. 5:**
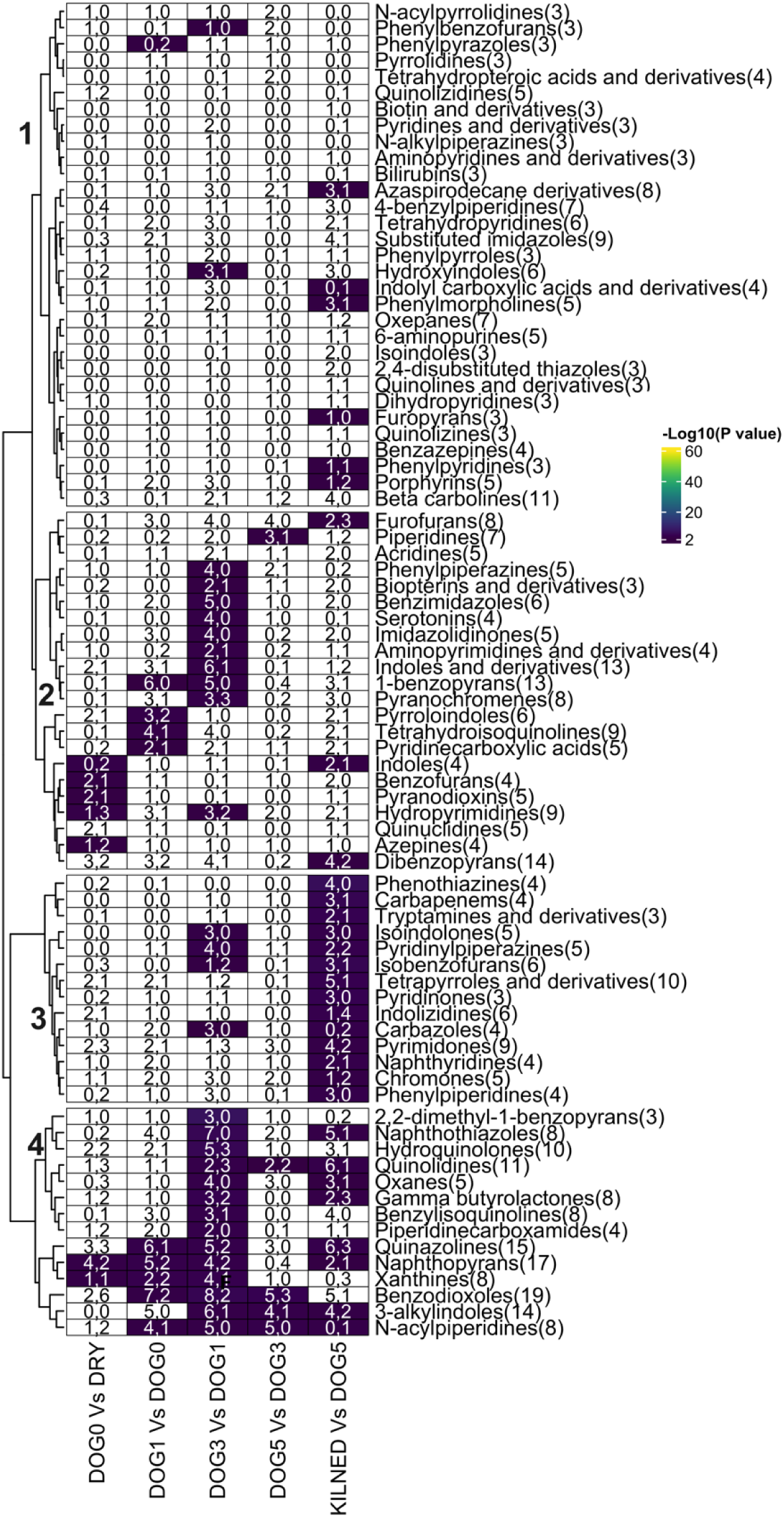
Hierarchical clustering analysis based on ChemRICH FDR in the five indicated stage comparisons of ClassyFire parent classes belonging to organoheterocyclic compound superclass. The clustering analysis was conducted using the Euclidean distance metric and the Ward.D2 clustering method. Color represents -log_10_(FDR) values. White indicates FDR > 0.01. Each cell contains two numbers, separated by a comma, corresponding to the number of metabolites in that parent class that significantly increased and that significantly decreased, respectively. The total number of metabolites detected in each parent class is indicated in parentheses.

Various OHCs derived from tryptamine were increased in malt relative to dry barley, such as N-palmitoyltryptamine (38-fold), N-arachidoyltryptamine (16-fold), 5-methoxytryptamine (1.2-fold), and N-desmethylzolmitriptan (17-fold) highlighting the importance of tryptophan metabolism during malting. Hydroxyindoles like N-acetyl serotonin (27-fold increase between DOG1 and DOG3, 25-fold in final malt) and 6-hydroxymelatonin (14-fold during kilning, 8.6-fold in final malt) also showed significant changes. Melatonin and its derivatives in beer have been reported to provide potential health benefits, such as antioxidant and anti-inflammatory properties (Maldonado et al., 2023) potentially making the final beer product more appealing to health-conscious consumers. However, benzenoids including styrenes, naphthalenes, anthracenes, and fluorenes, which predominantly increased during kilning (Table S9, Fig. S3B), negatively impact beer quality (Langos & Granvogl, 2016; Mastanjevíc et al., 2021).

Another compound class known for its pharmacological and biological roles is harmala alkaloids (Fig. S3C, cluster 1). Of the 11 detected harmala alkaloids, five accumulated in malt relative to dry barley (Table S9). Although there is limited information on harmala alkaloids in barley or malt, their presence has been previously reported in beer (Bosin & Faull, 1988). Therefore, harmala alkaloids may make their way into the final beverage via the malting process. Harmala alkaloids have been suggested to have health-related attributes (Moloudizargari et al., 2013). Effective management of malting conditions, particularly at the stages where these health-promoting or detrimental compounds are altered, can enhance the desirable sensory attributes and health benefits of the malt and eventually the finished beer.

##### 3.2.2.5 Lipid and lipid like molecules

The presence of lipids during the brewing process has both positive and negative influences on fermentation (Gordon et al., 2018). The current metabolome analysis revealed various lipid classes, categorized into five distinct clusters, showing dynamic changes during malting (Fig. S3D). The PC triterpenoids contained the highest number of lipid compounds (203), with most of these being heavily lipid-modified triterpenes (Table S7). Significant alterations in triterpenoid levels were observed during malting, with 80 compounds showing increased levels in the finished malt compared to the dry seed (Table S7). Triterpenoids, a subclass of terpenes, are produced by the cyclization of squalene and consist of six isoprene units. Various terpenic compounds have been reported in the volatile composition of a wide range of beer types (Hao et al., 2014; Martins et al., 2018; Riu-Aumatell et al., 2014; Tsuji & Mizuno, 2010). These compounds contribute to the aroma and flavor profile of beer. Different factors influencing the presence of terpenic compounds in beer have been studied, including the characteristics of hops and hopping regimes, fermentation processes, yeast metabolism (Dresel et al., 2015; King & Dickinson, 2003; Kollmannsberger et al., 2007; Praet et al., 2015). However, there is scant information available specifically on triterpenoids and their presence in malt. It would be interesting to explore the specific pathways through which triterpenoids are synthesized and modified during malting and how these modifications impact the final beer quality. Furthermore, it would be interesting to investigate whether these triterpenoids persist after malting and throughout the later stages of brewing.

### 3.3 Metabolic pathway and function analysis during malting

The previous analyses allowed us to identify specific chemical classes altered across different stages of malting. To better understand how these metabolome dynamics fit into metabolic processes, KEGG pathway enrichment analysis was performed using abundance data for metabolites with KEGG IDs (Table S10). 29 KEGG pathways showed significant changes (FDR < 0.05) in at least one transition between consecutive stages (Table S11, Fig. S4). A schematic representation of altered metabolic pathways is given in Fig. 6. The following sections discuss several altered pathways in detail, focusing on their specific functions and roles during malting.

**Fig. 6:**
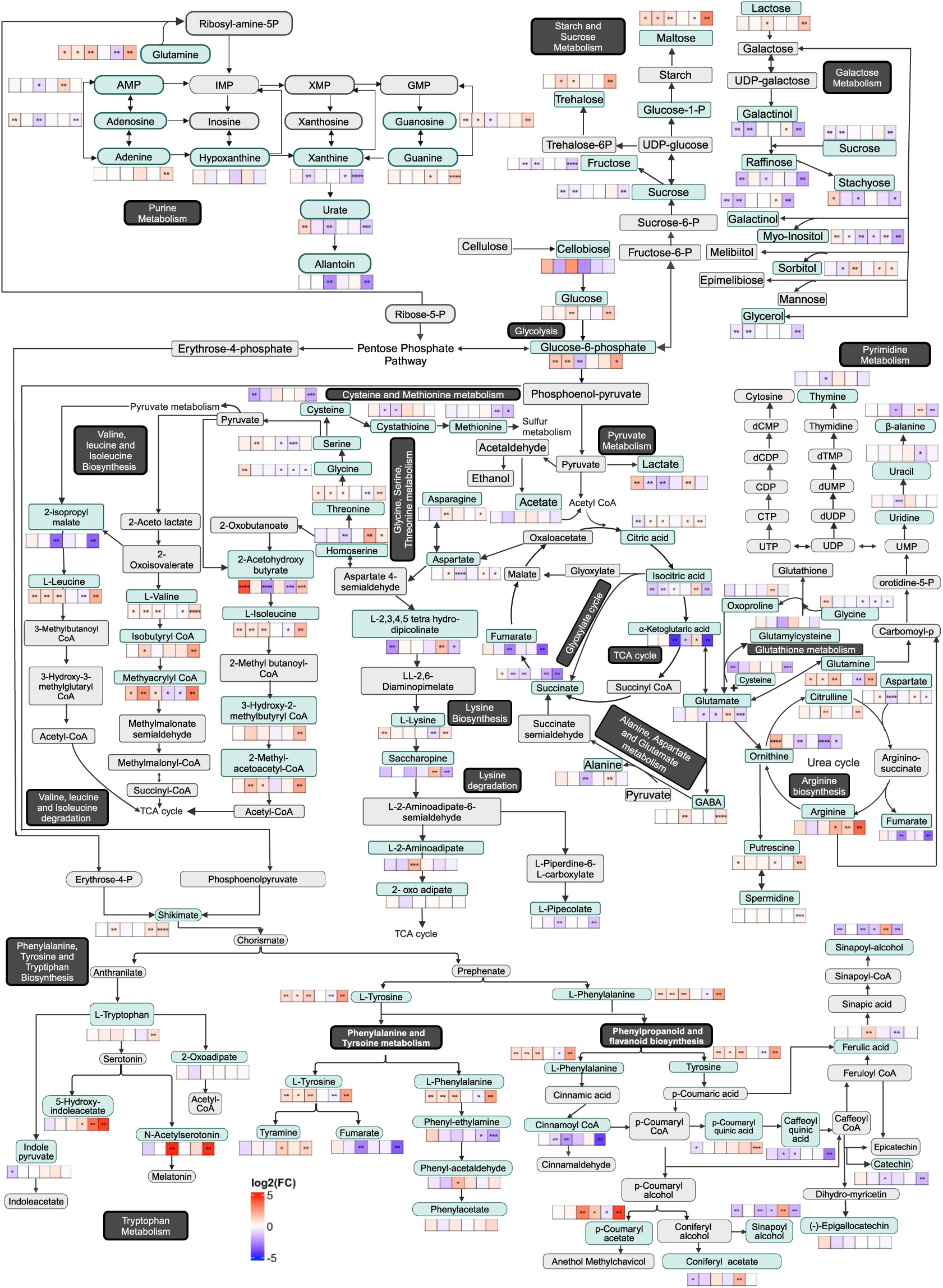
A schematic representation of KEGG metabolic pathways altered throughout the malting time course. Metabolites in green boxes were detected in our study, while those in grey boxes were not. Pathway names are indicated in black boxes. The individual heatmaps display the log2 fold change (FC) in the following six comparisons between malting stages, in this order: DOG0 Vs Dry, DOG1 Vs DOG0, DOG3 Vs DOG1, DOG5 Vs DOG3, Kilned Vs DOG5) and Kilned malt Vs Dry seed. Statistical significance is indicated by ‘*’ for FDR ≤ 0.05, ‘**’ for FDR ≤ 0.01, ‘***’ for FDR ≤ 0.001, and ‘****’ for FDR ≤ 0.0001. Note that the size or thickness of the arrows conveys no information regarding the chemical reactions.

#### 3.3.1 Dynamic energy metabolism and stress response during malting

Dry seeds are metabolically dormant, however upon imbibing water, metabolic activity resumes. The 2.5-fold increase in glucose-6-phosphate during steeping suggests activation of glycolysis for energy production (Fig. 6). A similar increase in glucose-6-phosphate has also been observed in imbibed wild oat seeds(Larondelle et al., 1987). Various proteins involved in catalyzing the conversion of glucose to pyruvate via glycolysis were reported in barley under hypoxia conditions (O’Lone et al., 2023). Glycolysis generates NADH and pyruvate, which, under normoxic conditions, are funneled into the TCA cycle via pyruvate dehydrogenase to produce ATP. Despite periodic air rests during steeping, grains face hypoxic conditions during immersion phases due to limited air diffusion. When comparing steeped grains to dry grains, we detected a significant increase in lactate, indicating that pyruvate is preferentially converted into lactate, thereby reducing TCA cycle activity and favoring anaerobic pathways for ATP synthesis under submerged conditions. This metabolic shift is further supported by the peak expression of genes associated with anaerobic respiration post-imbibition in soybean and barley (Bellieny-Rabelo et al., 2016; Vinje et al., 2021).

During steeping, the levels of TCA cycle intermediates such as citric and isocitric acid decreased, alpha-ketoglutaric acid levels remained stable, while succinate levels increased. The decrease in isocitrate and increase in succinate suggest the possible activation of the glyoxylate cycle. This aids in directly converting isocitrate to glyoxylate and succinate, bypassing two decarboxylation steps, potentially optimizing energy and carbon resource management under low-oxygen conditions that occurs during steeping. Activation of glyoxylate cycle under hypoxia has been documented in rice (Lu et al., 2005).

Gamma-aminobutyric acid (GABA), increased by 3.5-fold during steeping, correlating with upregulated expression of glutamate decarboxylase (HORVU.MOREX.r3.4HG0393690) during this phase in barley malting (Vinje et al., 2021). Under normoxia, GABA is catabolized via GABA transaminase to succinic semialdehyde, which in turn is oxidized via an NAD-dependent succinic semialdehyde dehydrogenase to succinate. The rise in GABA during steeping can be attributed to the inhibition of NAD-dependent succinic semialdehyde dehydrogenase (SSADH) under hypoxic conditions (Breitkreuz et al., 2003). A decrease in expression of SSADH was also reported during steeping in barley (Vinje et al., 2021). This increase in GABA is consistent with findings across various plant species exposed to low oxygen stress (Aurisano et al., 1995; Breitkreuz et al., 2003; Guo et al., 2012). As the grains transitioned from DOG0 to DOG3, GABA levels decreased by 1.7-fold, indicating a resumption of aerobic metabolism. During DOG3, in addition to decreases in GABA and lactate, there was a significant decline in key TCA cycle intermediates, such as a 21-fold decrease in alpha-ketoglutarate (Fig. 6). This decrease may reflect a rapid utilization of these intermediates for biosynthetic processes during early germination, as supported by the observed increase in various amino acids (Fig. S5) that are commonly synthesized from TCA cycle intermediates. Additionally, the expression of various TCA cycle enzymes was observed to increase in abundance after steeping, indicating a clear switch to aerobic respiration during germination (Vinje et al., 2021). During the kilning phase, there was an accumulation of early-stage TCA intermediates (Fig. S6A), suggesting alterations in metabolic rates or pathways rather than a cessation of metabolic activities. Consistent with earlier findings, GABA levels continued to decrease during kilning (Frank et al., 2011; Samaras et al., 2005).

A significant accumulation of ROS has been observed at the end of steeping as compared to dry seed (Mahalingam et al., 2021). Increased GABA concentration during steeping may serve as a potent antioxidant, when hypoxic conditions can disrupt the electron transport chain, leading to increased reactive oxygen species (ROS) generation and potential oxidative cell damage (Jacoby et al., 2018; O’Lone et al., 2023). Glutathione, another important antioxidant, was not detected by our analytical methods, but precursor compounds were identified. Glycine and gamma-glutamylcysteine both accumulated during steeping (1.8-fold and 1.25-fold, respectively), while glutamate and cysteine both decreased). These changes may indicate a preparatory phase for glutathione synthesis to counteract ROS produced during metabolic reactivation. Kilning appears to disrupt the gamma-glutamyl cycle, leading to oxoproline accumulation (1.8-fold). Oxoproline can accumulate under conditions of increased oxidative stress or disruption in glutathione metabolism. Oxoproline has also been observed to increase by 4-fold during kilning by Zhao et al., (2022). Furthermore, we observed a significant increase in the abundance of polyamines, such as putrescine and cadaverine during steeping (Fig. S6B). Polyamines have been shown to decrease hypoxia-induced oxidative damage by modulating the free radical scavenging system in various plant species (Reggiani et al., 1989; Wang et al., 2014; Yiu et al., 2009).

#### 3.3.2 Metabolic changes in starch and oligosaccharides metabolism during malting

During early malting, amylolytic enzymes target the primary storage reserve, starch, in the endosperm, converting it into simpler sugars essential for the germination process. In addition to starch, raffinose family oligosaccharides (RFOs) also serve as a readily metabolizable carbohydrate source for energy generation during germination. Moreover, RFOs regulate osmotic balance and help remove excess ROS, maintaining cell homeostasis (Nishizawa-Yokoi et al., 2008; Obendorf, 1997). Two of the RFOs, galactinol and raffinose, decreased significantly during steeping, while stachyose levels increased (Fig. 6). This suggests that during steeping, galactose from galactinol is added to sucrose to form raffinose, and stachyose synthase then produces stachyose from raffinose (Sengupta et al., 2015). Although stachyose was not previously detected in barley malting, it increased during early imbibition in adzuki beans (Peterbauer et al., 1999). Significant reductions in stachyose, sucrose, and raffinose levels by DOG3 indicate potential hydrolysis of these sugars, releasing galactose for glucose production, thereby supporting germination. RFOs are crucial respiratory substrates in germinating seeds, and their rapid breakdown is essential; disrupting this process can delay seed germination (Blöchl et al., 2007). Consistent with this study, raffinose also decreased during germination in barley (Zhao et al., 2022) and wheat (Byeon et al., 2022).

During steeping and DOG1, despite decreased sucrose levels, fructose did not rise correspondingly. This could be due to rapid fructose utilization for energy and growth. Monosaccharides and disaccharides (e.g., glucose, maltose, isomaltose, lactose, trehalose, xylose, β-genotobiose) increased significantly by the end of DOG3 (Table S5), while trisaccharides and tetrasaccharides (e.g., raffinose, stachyose) were reduced, suggesting active degradation of cell walls and conversion of stored polysaccharides into mono- and disaccharides to meet energy demands and mitigate stress during germination. The 6.6-fold increase in β-cellobiose between DOG1 and DOG3 further indicates active cell wall degradation (FDR = 0.35). Supporting this, transcript levels for various hydrolytic enzymes involved in cell wall degradation, starch depolymerization, and polysaccharide breakdown were found to increase in barley grains during early germination (Betts et al., 2017; Vinje et al., 2021). Additionally, hydrolytic enzyme activity increases during germination due to the release of bound inactive enzymes by proteolytic activity (Betts et al., 2017). As the energy demands reduce during later stages of growth (DOG3 to DOG5) most sugars do not exhibit significant changes. Tagatose, a valuable sugar in the current market because of its lower caloric value compared to sucrose and approximately 90% of the sweetness of sucrose (Ravikumar et al., 2021), was observed to increase two-fold during kilning (Fig. S6A). Future research should precisely quantify tagatose levels and biochemical processes responsible for its increase during kilning, aiming to produce low calorie beer, yet still retaining a desirable level of sweetness through optimized malting.

#### 3.3.3 Metabolic changes in amino acid metabolism during malting

The changes during malting of the 20 proteinaceous amino acids are illustrated in Fig. S5. Levels of most amino acids, including branched-chain amino acids (BCAAs) and phenylalanine, which are important for beer flavor (Kathuria et al., 2024), exhibited increased levels in kilned malt than in the starting dry grain. However, five amino acids (cysteine, methionine, glycine, aspartic and glutamic acid) decreased by the end of malting. The reduction in sulfur-containing amino acids (cysteine and methionine) was particularly strong and can be attributed to their role as precursors of volatile sulfur compounds, which influence the sensory quality of beer (Mikulíková et al., 2009). During kilning, methionine is particularly susceptible to degradation through interactions with reducing sugars, leading to the formation of dimethyl sulfide and methional via the Strecker degradation process. High concentrations of dimethyl sulfide impart an undesirable cooked sweetcorn flavor to beer (Bogdan & Kordialik-Bogacka, 2017). Furthermore, when temperatures exceed 60°C, S-methylmethionine (a derivative of methionine) breaks down into homoserine and additional dimethyl sulfide, evidenced by a 4-fold increase in homoserine during kilning (Mikulíková et al., 2009) (Fig. S6C). The overall decrease in aspartate and glutamate occurred mainly during germination when they may be converted to other proteinaceous amino acids for protein synthesis (Fig. S5) and catabolized to intermediates for the TCA cycle (Fig. 6).

A significant increase in BCAAs is observed progressively until DOG3, followed by a marked decrease in their synthesis. However, the expected rise in BCAA levels at DOG3 was not as high, due to their rapid catabolism and utilization in energy metabolism pathways, as indicated by the increase in intermediates of BCAA catabolism (Fig. 6). During kilning, there is a significant decline in BCAAs, likely due to their conversion into Strecker aldehydes; valine to 2-methylpropanal, isoleucine to 2-methylbutanal, and leucine to 3-methylbutanal (Filipowska et al., 2021). Despite the decrease during kilning, BCAAs demonstrated increased abundance in the final malt compared to dry seed. This increased abundance of BCAAs in malt promotes the formation of higher alcohols and esters by yeast through the Ehrlich amino acid degradation pathway, playing a crucial role in defining the distinctive characteristics of beer (Lin et al., 2022). Additionally, shikimic acid exhibited a significant increase at DOG1 likely due to the activation of the shikimate pathway, which is crucial for the biosynthesis of aromatic amino acids (AAs). AAs followed a similar trend throughout the malting stages, though changes in tryptophan levels are statistically non-significant. Tryptophan appears to undergo multiple degradation pathways (Fig. 6). Most amino acids showed a decrease during kilning, possibly due to their involvement in the Maillard reaction.

## 4. Conclusion

While previous research has extensively examined specific biochemical transformations and the roles of storage carbohydrates and proteins during malting, a more comprehensive understanding of metabolite dynamics across malting stages has been less explored. Our integrative approach, employing both GC-MS and LC-MS, provides detailed snapshots of the metabolic landscape during malting. This dataset may serve as a benchmark for assessing how variations in genotype and malting protocols influence metabolic transformations during malting and the quality of the final malt. Furthermore, it may facilitate future research aimed at enhancing specific metabolic pathways and desirable metabolite traits in the final malt. Among our novel findings, we identified a diverse array of microbial metabolites, many of which were elevated in the final malt. This underscores the need for future research into the dynamic and integral role of the microbial ecosystem in malting, moving beyond merely focusing on specific fungal species or toxins. Additionally, the identification of beneficial microbial metabolites and the cognate microbe would offer opportunities to develop novel traits for microbial quality assessments, ensuring the production of high-quality, safe malted barley.

## Acknowledegements

GC-MS metabolomics data were acquired and processed at the UC Davis West Coast Metabolomics Center. LC-MS metabolomics data were acquired and processed by Corey Broeckling and colleagues in the Colorado State University Analytical Resources Core (RRID: SCR_021758). The micro-malting and malt quality analysis were performed at the USDA-ARS Cereal Crops Research Unit, Madison WI by Dr. Marcus Vinje, Michael O’Connor, Bryan Lemmenes, and Chris Martens. Figure 6 was created with BioRender (https://app.biorender.com/).

## CRediT authorship contribution statement

**Heena Rani:** Writing – review & editing, Writing – original draft, Visualization, Methodology, Investigation, Formal analysis, Data curation, Conceptualization. **Sarah J Whitcomb:** Writing – review & editing, Supervision, Resources, Project administration, Methodology, Investigation, Formal analysis, Data curation, Conceptualization.

## Abbreviations

AA: Aromatic amino acid
ACN: Acetonitrile
AMBA: American Malting Barley Association
ANOVA: Analysis of Variance
ATP: Adenosine Triphosphate
BCAA: Branched-Chain Amino Acid
ChemRICH: Chemical similarity enrichment analysis
CDP: Cyclic Depsipeptide
DOG: Days of Germination
FA: Fatty Acid
FAME: Fatty Acid Methyl Ester
FDR: False Discovery Rate
GABA: Gamma-aminobutyric acid
GC-MS: Gas Chromatography-Mass Spectrometry
GTP: Guanosine Triphosphate
HCA: Hierarchical Clustering Analysis
IPA: Isopropanol
KEGG: Kyoto Encyclopedia of Genes and Genomes
LC-MS: Liquid Chromatography-Mass Spectrometry
MS: Mass Spectrometry
MSE: MSTFA: N-methyl-N-trimethylsilyltrifluoroacetamide
NMR: Nuclear Magnetic Resonance
OHC: Organo-Heterocyclic Compound
OOC: Organo Oxygen Compounds
OAD: Organic Acid and Derivatives
PA: Polyamine
PCA: Principal Component Analysis
PC: Principal Component
PC: Parent Class
PP: Phenylpropanoid and Polyketides
RFO: Raffinose Family Oligosaccharides
QC: Quality Control
ROS: Reactive Oxygen Species
TOF: Time of Flight
TQMS: Triple Quadrupole Mass Spectrometry
UPLC: Ultra-Performance Liquid Chromatography
TCA: Tricarboxylic Acid
ZEN: Zearalenone

