## Supplementary Fig S2-S6 for "Integrative LC-MS and GC-MS Metabolic Profiling Unveils Dynamic Changes during Barley Malting"

**Supplementary Figures**

**Fig. S1**: An interactive pie chart showing the classification and distribution of annotated and identified metabolites at different levels—superclass, class, subclass, and parent class. The pie chart on the left represents data from LC-MS, and the pie chart on the right represents data from GC-MS.

**Fig. S2**: Scores plot of principal component analysis after autoscaling (A) LC-MS dataset comprising 8921 metabolites and (B) GC-MS dataset comprising 773 metabolites. Different symbols indicate the three distinct malting batches, each with three replicates, while colors represent the malting stage and pooled quality control sample.

**Fig. S3**: Hierarchical clustering analysis of ClassyFire parent classes based on ChemRICH FDR in the five indicated stage comparisons. Clustering analysis was conducted using the Euclidean distance metric and the Ward.D2 clustering method. Color represents $-\log_{10}$(FDR) values. White indicates FDR > 0.01. Each cell contains two numbers, separated by a comma, corresponding to the number of metabolites in that parent class that were significantly increased and significantly decreased, respectively. The total number of metabolites detected in each parent class is indicated in parentheses. (A) Parent classes belonging to lignans, neolignans, and related compounds superclass, (B) Parent classes belonging to benzenoids superclass, (C) Parent classes belonging to alkaloids and derivatives superclass, (D) Parent classes belonging to lipids and lipid-like molecules superclass.

**Fig. S4**: Hierarchical clustering analysis of pathways (with five or more hits) in the five indicated stage comparisons. The analysis was conducted using the Euclidean distance metric and the Ward.D2 clustering method. The color scale represents -log10(FDR) values, with white indicating an FDR > 0.05. Numbers in parentheses next to pathway names denote the total number of compounds in the pathway, the number of compounds matched (hits) from the user-uploaded data, and the number of significant hits based on ANOVA analysis across all stages (FDR < 0.01).

**Fig. S5**: Variations in the abundance of 20 key amino acids in the six indicated comparisons. Color indicates the log2(fold change). Statistical significance is indicated by '*' for FDR ≤ 0.05, '**' for FDR ≤ 0.01, '***' for FDR ≤ 0.001, and '****' for FDR ≤ 0.0001.

**Fig. S6**: Heatmaps displaying metabolites with known KEGG IDs, categorized into super pathways (A) Carbohydrate Metabolism, (B) Metabolism of Other Amino Acids, (C) Amino Acid Metabolism, (D) Biosynthesis of Other Secondary Metabolites, (E) Lipid Metabolism, and (F) Nucleotide Metabolism. Metabolites are arranged based on sub-pathways within each super pathway. The individual heatmaps display the log2 fold change (FC) in the indicated six comparisons. Statistical significance is indicated by '*' for FDR ≤ 0.05, '**' for FDR ≤ 0.01, '***' for FDR ≤ 0.001, and '****' for FDR ≤ 0.0001.

### Fig S2

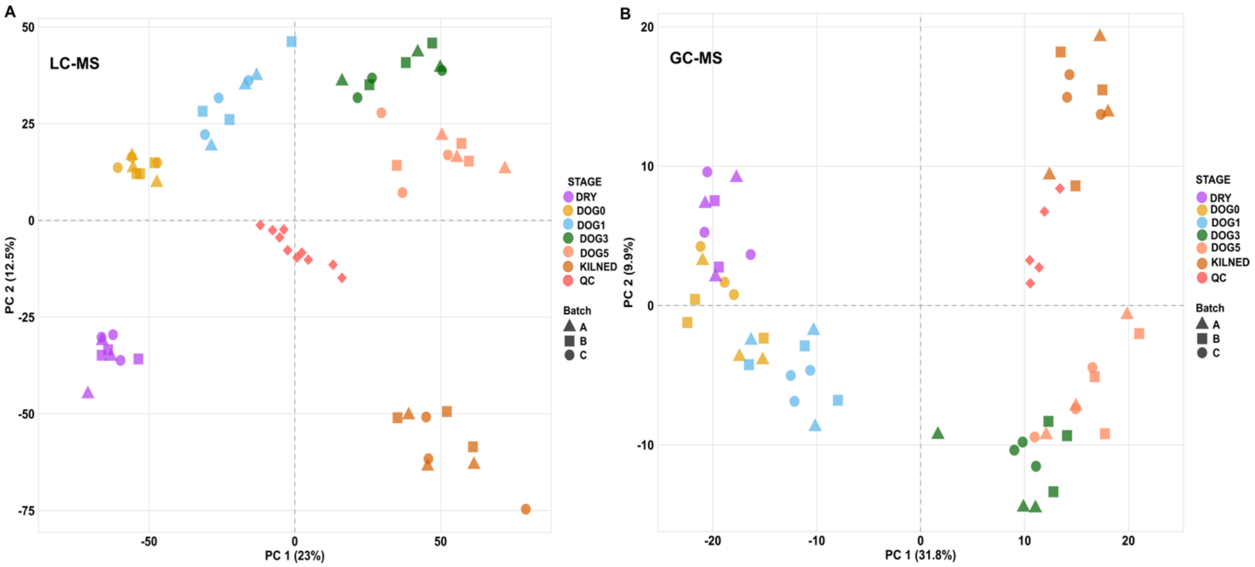
A
LC-MS
PC 1 (23%)
PC 2 (12.5%)
STAGE
DRY
DOG0
DOG1
DOG3
DOG5
KILNED
QC
Batch
A
B
C
B
GC-MS
PC 1 (31.8%)
PC 2 (9.9%)
STAGE
DRY
DOG0
DOG1
DOG3
DOG5
KILNED
QC
Batch
A
B
C

**Fig S3**

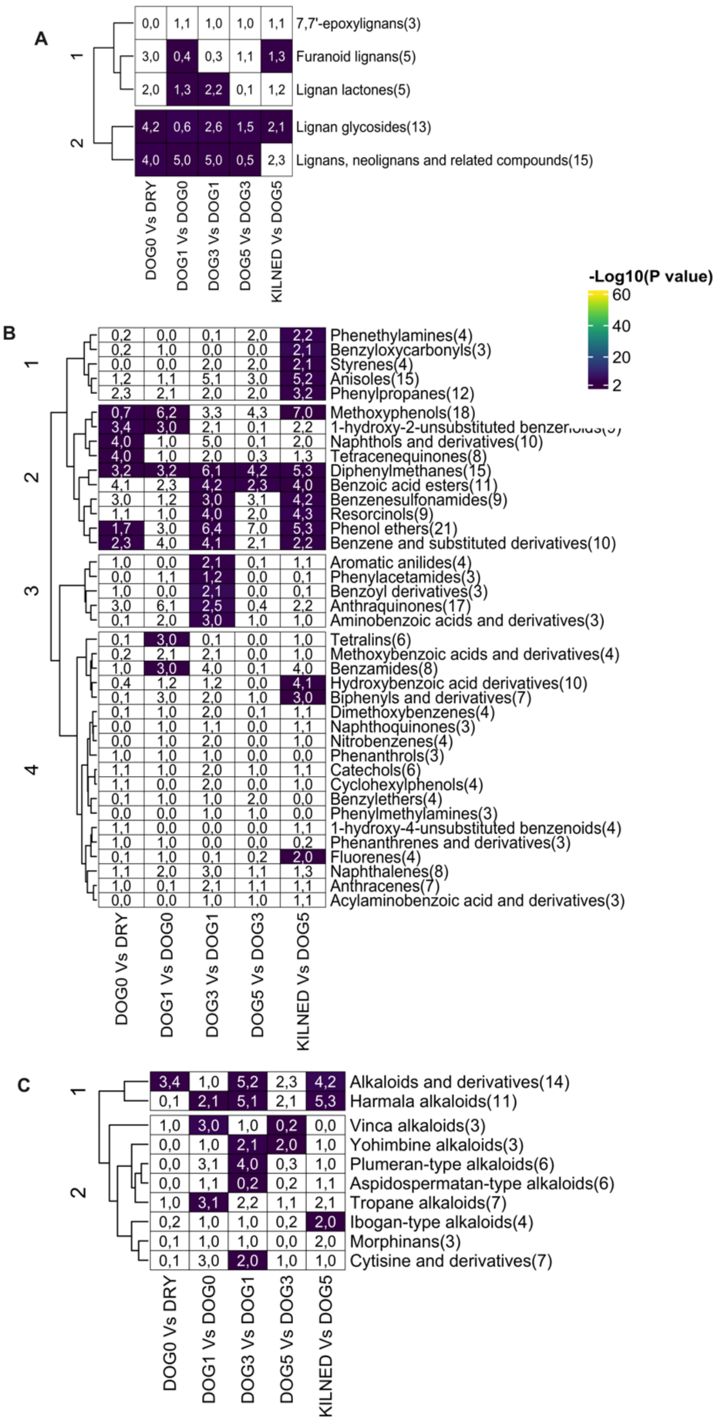
A
1
2
0,0 1,1 1,0 1,0 1,1 7,7'-epoxylignans(3)
3,0 0,4 0,3 1,1 1,3 Furanoid lignans(5)
2,0 1,3 2,2 0,1 1,2 Lignan lactones(5)
4,2 0,6 2,6 1,5 2,1 Lignan glycosides(13)
4,0 5,0 5,0 0,5 2,3 Lignans, neolignans and related compounds(15)
DOG0 Vs DRY
DOG1 Vs DOG0
DOG3 Vs DOG1
DOG5 Vs DOG3
KILNED Vs DOG5
-Log10(P value)
60
40
20
2
B
1
2
3
4
0,2 0,0 0,1 2,0 2,2 Phenethylamines(4)
0,2 1,0 0,0 0,0 2,1 Benzyloxycarbonyls(3)
0,0 0,0 2,0 2,0 2,1 Styrenes(4)
1,2 1,1 5,1 3,0 5,2 Anisoles(15)
2,3 2,1 2,0 2,0 3,2 Phenylpropanes(12)
0,7 6,2 3,3 4,3 7,0 Methoxyphenols(18)
3,4 3,0 2,1 0,1 2,2 1-hydroxy-2-unsubstituted benzen
4,0 1,0 5,0 0,1 2,0 Naphthols and derivatives(10)
4,0 1,0 2,0 0,3 1,3 Tetracenequinones(8)
3,2 3,2 6,1 4,2 5,3 Diphenylmethanes(15)
4,1 2,3 4,2 2,3 4,0 Benzoic acid esters(11)
3,0 1,2 3,0 3,1 4,2 Benzenesulfonamides(9)
1,1 1,0 4,0 2,0 4,3 Resorcinols(9)
1,7 3,0 6,4 7,0 5,3 Phenol ethers(21)
2,3 4,0 4,1 2,1 2,2 Benzene and substituted derivatives(10)
1,0 0,0 2,1 0,1 1,1 Aromatic anilides(4)
0,0 1,1 1,2 0,0 0,1 Phenylacetamides(3)
1,0 0,0 2,1 0,0 0,1 Benzoyl derivatives(3)
3,0 6,1 2,5 0,4 2,2 Anthraquinones(17)
0,1 2,0 3,0 1,0 1,0 Aminobenzoic acids and derivatives(3)
0,1 3,0 0,1 0,0 1,0 Tetralins(6)
0,2 2,1 2,1 0,0 1,0 Methoxybenzoic acids and derivatives(4)
1,0 3,0 4,0 0,1 4,0 Benzamides(8)
0,4 1,2 1,2 0,0 4,1 Hydroxybenzoic acid derivatives(10)
0,1 3,0 2,0 1,0 3,0 Biphenyls and derivatives(7)
0,1 1,0 2,0 0,1 1,1 Dimethoxybenzenes(4)
0,0 1,0 1,1 0,0 1,1 Naphthoquinones(3)
0,0 1,0 2,0 0,0 1,0 Nitrobenzenes(4)
1,0 1,0 1,0 0,0 0,0 Phenanthrols(3)
1,1 1,0 2,0 1,0 1,1 Catechols(6)
1,1 0,0 2,0 0,0 1,0 Cyclohexylphenols(4)
0,1 1,0 1,0 2,0 0,0 Benzylethers(4)
0,0 0,0 1,0 1,0 0,0 Phenylmethylamines(3)
1,1 0,0 0,0 0,0 1,1 1-hydroxy-4-unsubstituted benzenoids(4)
1,0 1,0 0,0 0,0 0,2 Phenanthrenes and derivatives(3)
0,1 1,0 0,1 0,2 2,0 Fluorenes(4)
1,1 2,0 3,0 1,1 1,3 Naphthalenes(8)
1,0 0,1 2,1 1,1 1,1 Anthracenes(7)
0,0 0,0 1,0 1,0 1,1 Acylaminobenzoic acid and derivatives(3)
DOG0 Vs DRY
DOG1 Vs DOG0
DOG3 Vs DOG1
DOG5 Vs DOG3
KILNED Vs DOG5
C
1
2
3,4 1,0 5,2 2,3 4,2 Alkaloids and derivatives(14)
0,1 2,1 5,1 2,1 5,3 Harmala alkaloids(11)
1,0 3,0 1,0 0,2 0,0 Vinca alkaloids(3)
0,0 1,0 2,1 2,0 1,0 Yohimbine alkaloids(3)
0,0 3,1 4,0 0,3 1,0 Plumeran-type alkaloids(6)
0,0 1,1 0,2 0,2 1,1 Aspidospermatan-type alkaloids(6)
1,0 3,1 2,2 1,1 2,1 Tropane alkaloids(7)
0,2 1,0 1,0 0,2 2,0 Ibogan-type alkaloids(4)
0,1 1,0 1,0 0,0 2,0 Morphinans(3)
0,1 3,0 2,0 1,0 1,0 Cytisine and derivatives(7)
DOG0 Vs DRY
DOG1 Vs DOG0
DOG3 Vs DOG1
DOG5 Vs DOG3
KILNED Vs DOG5

D

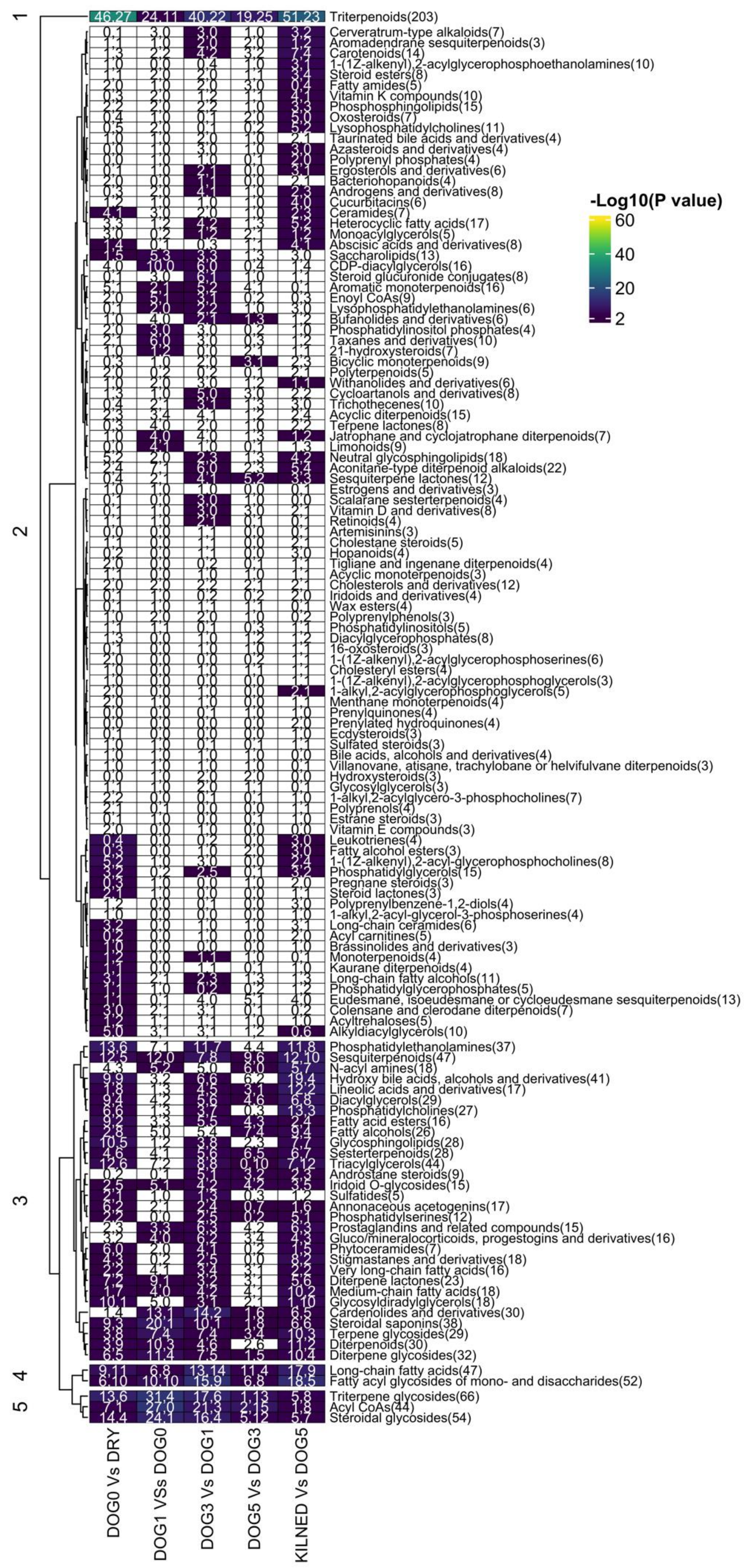

-Log10(P value)
60
40
20
2
1
2
3
4
5
Triterpenoids(203)
Cerveratrum-type alkaloids(7)
Aromadendrane sesquiterpenoids(3)
Carotenoids(14)
1-(1Z-alkenyl),2-acylglycerophosphoethanolamines(10)
Steroid esters(8)
Fatty amides(5)
Vitamin K compounds(10)
Phosphosphingolipids(15)
Oxosteroids(7)
Lysophosphatidylcholines(11)
Taurinated bile acids and derivatives(4)
Azasteroids and derivatives(4)
Polyprenyl phosphates(4)
Ergosterols and derivatives(6)
Bacteriohopanoids(4)
Androgens and derivatives(8)
Cucurbitacins(6)
Ceramides(7)
Heterocyclic fatty acids(17)
Monoacylglycerols(5)
Abscisic acids and derivatives(8)
Saccharolipids(13)
CDP-diacylglycerols(16)
Steroid glucuronide conjugates(8)
Aromatic monoterpenoids(16)
Enoyl CoAs(9)
Lysophosphatidylethanolamines(6)
Bufanolides and derivatives(6)
Phosphatidylinositol phosphates(4)
Taxanes and derivatives(10)
21-hydroxysteroids(7)
Bicyclic monoterpenoids(9)
Polyterpenoids(5)
Withanolides and derivatives(6)
Cycloartanols and derivatives(8)
Trichothecenes(10)
Acyclic diterpenoids(15)
Terpene lactones(8)
Jatrophane and cyclojatrophane diterpenoids(7)
Limonoids(9)
Neutral glycosphingolipids(18)
Aconitane-type diterpenoid alkaloids(22)
Sesquiterpene lactones(12)
Estrogens and derivatives(3)
Scalarane sesterterpenoids(4)
Vitamin D and derivatives(8)
Retinoids(4)
Artemisinins(3)
Cholestane steroids(5)
Hopanoids(4)
Tigliane and ingenane diterpenoids(4)
Acyclic monoterpenoids(3)
Cholesterols and derivatives(12)
Iridoids and derivatives(4)
Wax esters(4)
Polyprenylphenols(3)
Phosphatidylinositols(5)
Diacylglycerophosphates(8)
16-oxosteroids(3)
1-(1Z-alkenyl),2-acylglycerophosphoserines(6)
Cholesteryl esters(4)
1-(1Z-alkenyl),2-acylglycerophosphoglycerols(3)
1-alkyl,2-acylglycerophosphoglycerols(5)
Menthane monoterpenoids(4)
Prenylquinones(4)
Prenylated hydroquinones(4)
Ecdysteroids(3)
Sulfated steroids(3)
Bile acids, alcohols and derivatives(4)
Villanovane, atisane, trachylobane or helvifulvane diterpenoids(3)
Hydroxysteroids(3)
Glycosylglycerols(3)
1-alkyl,2-acylglycero-3-phosphocholines(7)
Polyprenols(4)
Estrane steroids(3)
Vitamin E compounds(3)
Leukotrienes(4)
Fatty alcohol esters(3)
1-(1Z-alkenyl),2-acyl-glycerophosphocholines(8)
Phosphatidylglycerols(15)
Pregnane steroids(3)
Steroid lactones(3)
Polyprenylbenzene-1,2-diols(4)
1-alkyl,2-acyl-glycerol-3-phosphoserines(4)
Long-chain ceramides(6)
Acyl carnitines(5)
Brassinolides and derivatives(3)
Monoterpenoids(4)
Kaurane diterpenoids(4)
Long-chain fatty alcohols(11)
Phosphatidylglycerophosphates(5)
Eudesmane, isoeudesmane or cycloeudesmane sesquiterpenoids(13)
Colensane and clerodane diterpenoids(7)
Acyltrehaloses(5)
Alkyldiacylglycerols(10)
Phosphatidylethanolamines(37)
Sesquiterpenoids(47)
N-acyl amines(18)
Hydroxy bile acids, alcohols and derivatives(41)
Lineolic acids and derivatives(17)
Diacylglycerols(29)
Phosphatidylcholines(27)
Fatty acid esters(16)
Fatty alcohols(26)
Glycosphingolipids(28)
Sesterterpenoids(28)
Triacylglycerols(44)
Androstane steroids(9)
Iridoid O-glycosides(15)
Sulfatides(5)
Annonaceous acetogenins(17)
Phosphatidylserines(12)
Prostaglandins and related compounds(15)
Gluco/mineralocorticoids, progestogins and derivatives(16)
Phytoceramides(7)
Stigmastanes and derivatives(18)
Very long-chain fatty acids(16)
Diterpene lactones(23)
Medium-chain fatty acids(18)
Glycosyldiradylglycerols(18)
Cardenolides and derivatives(30)
Steroidal saponins(38)
Terpene glycosides(29)
Diterpenoids(30)
Diterpene glycosides(32)
Long-chain fatty acids(47)
Fatty acyl glycosides of mono- and disaccharides(52)
Triterpene glycosides(66)
Acyl CoAs(44)
Steroidal glycosides(54)
DOG0 Vs DRY
DOG1 VSs DOG0
DOG3 Vs DOG1
DOG5 Vs DOG3
KILNED Vs DOG5

**Fig S4**

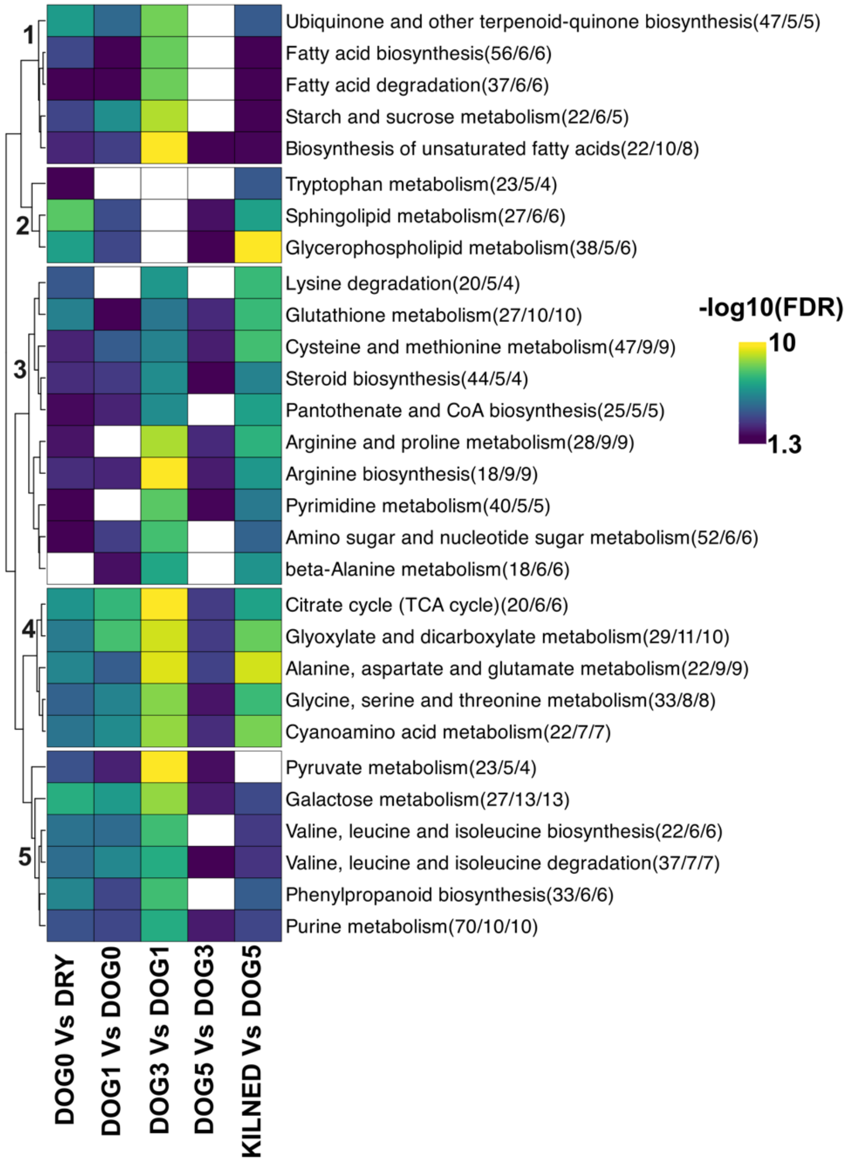
1
2
3
4
5
Ubiquinone and other terpenoid-quinone biosynthesis(47/5/5)
Fatty acid biosynthesis(56/6/6)
Fatty acid degradation(37/6/6)
Starch and sucrose metabolism(22/6/5)
Biosynthesis of unsaturated fatty acids(22/10/8)
Tryptophan metabolism(23/5/4)
Sphingolipid metabolism(27/6/6)
Glycerophospholipid metabolism(38/5/6)
Lysine degradation(20/5/4)
Glutathione metabolism(27/10/10)
Cysteine and methionine metabolism(47/9/9)
Steroid biosynthesis(44/5/4)
Pantothenate and CoA biosynthesis(25/5/5)
Arginine and proline metabolism(28/9/9)
Arginine biosynthesis(18/9/9)
Pyrimidine metabolism(40/5/5)
Amino sugar and nucleotide sugar metabolism(52/6/6)
beta-Alanine metabolism(18/6/6)
Citrate cycle (TCA cycle)(20/6/6)
Glyoxylate and dicarboxylate metabolism(29/11/10)
Alanine, aspartate and glutamate metabolism(22/9/9)
Glycine, serine and threonine metabolism(33/8/8)
Cyanoamino acid metabolism(22/7/7)
Pyruvate metabolism(23/5/4)
Galactose metabolism(27/13/13)
Valine, leucine and isoleucine biosynthesis(22/6/6)
Valine, leucine and isoleucine degradation(37/7/7)
Phenylpropanoid biosynthesis(33/6/6)
Purine metabolism(70/10/10)
DOG0 Vs DRY
DOG1 Vs DOG0
DOG3 Vs DOG1
DOG5 Vs DOG3
KILNED Vs DOG5
-log10(FDR)
10
1.3

**Fig S5**

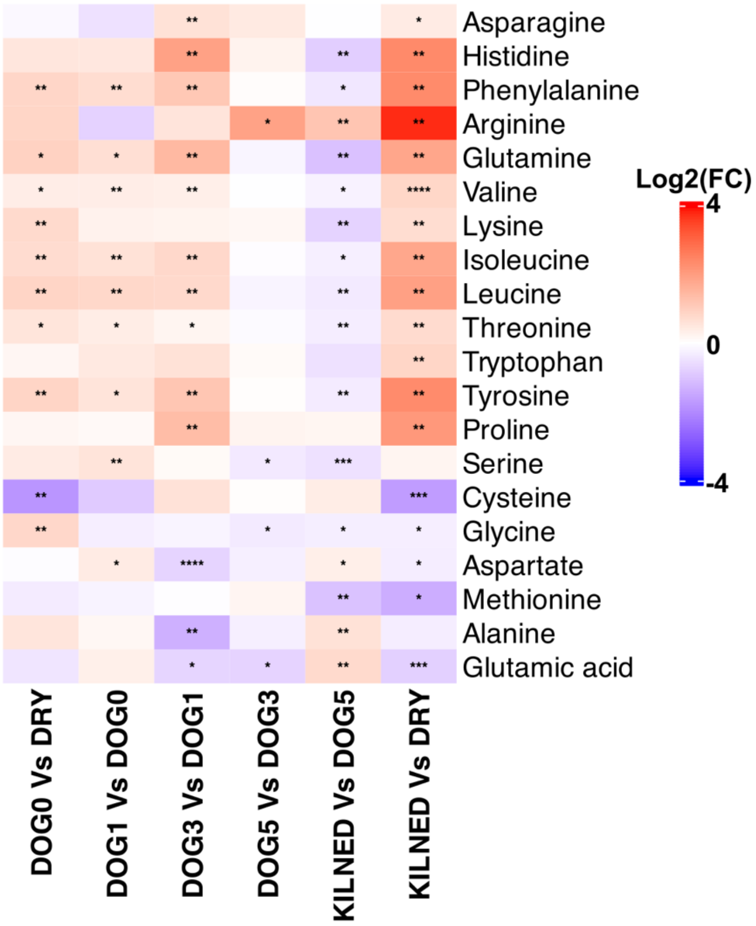
Asparagine
Histidine
Phenylalanine
Arginine
Glutamine
Valine
Lysine
Isoleucine
Leucine
Threonine
Tryptophan
Tyrosine
Proline
Serine
Cysteine
Glycine
Aspartate
Methionine
Alanine
Glutamic acid
DOG0 Vs DRY
DOG1 Vs DOG0
DOG3 Vs DOG1
DOG5 Vs DOG3
KILNED Vs DOG5
KILNED Vs DRY
Log2(FC)
4
0
-4

### Fig S6

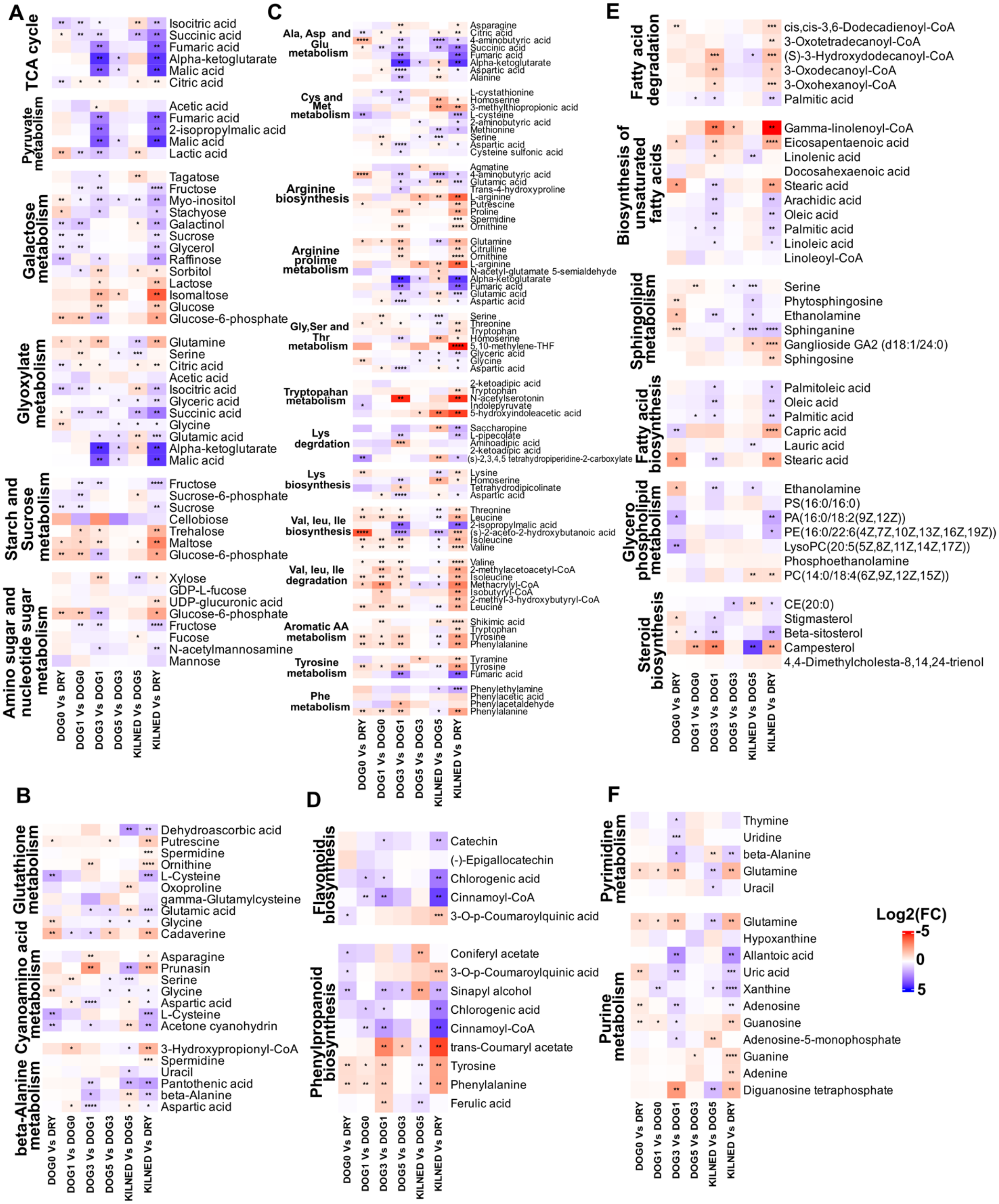
A
TCA cycle
Isocitric acid
Succinic acid
Fumaric acid
Alpha-ketoglutarate
Malic acid
Citric acid
Pyruvate metabolism
Acetic acid
Fumaric acid
2-isopropylmalic acid
Malic acid
Lactic acid
Galactose metabolism
Tagatose
Fructose
Myo-inositol
Stachyose
Galactinol
Sucrose
Glycerol
Raffinose
Sorbitol
Lactose
Isomaltose
Glucose
Glucose-6-phosphate
Glyoxylate metabolism
Glutamine
Serine
Citric acid
Acetic acid
Isocitric acid
Glyceric acid
Succinic acid
Glycine
Glutamic acid
Alpha-ketoglutarate
Malic acid
Starch and Sucrose metabolism
Fructose
Sucrose-6-phosphate
Sucrose
Cellobiose
Trehalose
Maltose
Glucose-6-phosphate
Amino sugar and nucleotide sugar metabolism
Xylose
GDP-L-fucose
UDP-glucuronic acid
Glucose-6-phosphate
Fructose
Fucose
N-acetylmannosamine
Mannose
DOG0 Vs DRY
DOG1 Vs DOG0
DOG3 Vs DOG1
DOG5 Vs DOG3
KILNED Vs DOG5
KILNED Vs DRY
C
Ala, Asp and Glu metabolism
Asparagine
Citric acid
4-aminobutyric acid
Succinic acid
Fumaric acid
Alpha-ketoglutarate
Aspartic acid
Alanine
Cys and Met metabolism
L-cystathionine
Homoserine
3-methylthiopropionic acid
L-cysteine
2-aminobutyric acid
Methionine
Serine
Aspartic acid
Cysteine sulfonic acid
Arginine biosynthesis
Agmatine
4-aminobutyric acid
Glutamic acid
Trans-4-hydroxyproline
L-arginine
Putrescine
Proline
Spermidine
Ornithine
Arginine proline metabolism
Glutamine
Citrulline
Ornithine
L-arginine
N-acetyl-glutamate 5-semialdehyde
Alpha-ketoglutarate
Fumaric acid
Glutamic acid
Aspartic acid
Gly,Ser and Thr metabolism
Serine
Threonine
Tryptophan
Homoserine
5,10-methylene-THF
Glyceric acid
Glycine
Aspartic acid
Tryptophan metabolism
2-ketoadipic acid
Tryptophan
N-acetylserotonin
Indolepyruvate
5-hydroxyindoleacetic acid
Lys degrdation
Saccharopine
L-pipecolate
Aminoadipic acid
2-ketoadipic acid
(s)-2,3,4,5 tetrahydropiperidine-2-carboxylate
Lys biosynthesis
Lysine
Homoserine
Tetrahydrodipicolinate
Aspartic acid
Val, leu, Ile biosynthesis
Threonine
Leucine
2-isopropylmalic acid
(s)-2-aceto-2-hydroxybutanoic acid
Isoleucine
Valine
Val, leu, Ile degradation
Valine
2-methylacetoacetyl-CoA
Isoleucine
Methacrylyl-CoA
Isobutyryl-CoA
2-methyl-3-hydroxybutyryl-CoA
Leucine
Aromatic AA metabolism
Shikimic acid
Tryptophan
Tyrosine
Phenylalanine
Tyrosine metabolism
Tyramine
Tyrosine
Fumaric acid
Phe metabolism
Phenylethylamine
Phenylacetic acid
Phenylacetaldehyde
Phenylalanine
DOG0 Vs DRY
DOG1 Vs DOG0
DOG3 Vs DOG1
DOG5 Vs DOG3
KILNED Vs DOG5
KILNED Vs DRY
E
Fatty acid degradation
cis,cis-3,6-Dodecadienoyl-CoA
3-Oxotetradecanoyl-CoA
(S)-3-Hydroxydodecanoyl-CoA
3-Oxodecanoyl-CoA
3-Oxohexanoyl-CoA
Palmitic acid
Biosynthesis of unsaturated fatty acids
Gamma-linolenoyl-CoA
Eicosapentaenoic acid
Linolenic acid
Docosahexaenoic acid
Stearic acid
Arachidic acid
Oleic acid
Palmitic acid
Linoleic acid
Linoleoyl-CoA
Sphingolipid metabolism
Serine
Phytosphingosine
Ethanolamine
Sphinganine
Ganglioside GA2 (d18:1/24:0)
Sphingosine
Fatty acid biosynthesis
Palmitoleic acid
Oleic acid
Palmitic acid
Capric acid
Lauric acid
Stearic acid
Glycero phospholipid metabolism
Ethanolamine
PS(16:0/16:0)
PA(16:0/18:2(9Z,12Z))
PE(16:0/22:6(4Z,7Z,10Z,13Z,16Z,19Z))
LysoPC(20:5(5Z,8Z,11Z,14Z,17Z))
Phosphoethanolamine
PC(14:0/18:4(6Z,9Z,12Z,15Z))
Steroid biosynthesis
CE(20:0)
Stigmasterol
Beta-sitosterol
Campesterol
4,4-Dimethylcholesta-8,14,24-trienol
DOG0 Vs DRY
DOG1 Vs DOG0
DOG3 Vs DOG1
DOG5 Vs DOG3
KILNED Vs DOG5
KILNED Vs DRY
B
Glutathione metabolism
Dehydroascorbic acid
Putrescine
Spermidine
Ornithine
L-Cysteine
Oxoproline
gamma-Glutamylcysteine
Glutamic acid
Glycine
Cadaverine
Cyanoamino acid metabolism
Asparagine
Prunasin
Serine
Glycine
Aspartic acid
L-Cysteine
Acetone cyanohydrin
beta-Alanine metabolism
3-Hydroxypropionyl-CoA
Spermidine
Uracil
Pantothenic acid
beta-Alanine
Aspartic acid
DOG0 Vs DRY
DOG1 Vs DOG0
DOG3 Vs DOG1
DOG5 Vs DOG3
KILNED Vs DOG5
KILNED Vs DRY
D
Flavonoid biosynthesis
Catechin
(-)-Epigallocatechin
Chlorogenic acid
Cinnamoyl-CoA
3-O-p-Coumaroylquinic acid
Phenylpropanoid biosynthesis
Coniferyl acetate
3-O-p-Coumaroylquinic acid
Sinapyl alcohol
Chlorogenic acid
Cinnamoyl-CoA
trans-Coumaryl acetate
Tyrosine
Phenylalanine
Ferulic acid
DOG0 Vs DRY
DOG1 Vs DOG0
DOG3 Vs DOG1
DOG5 Vs DOG3
KILNED Vs DOG5
KILNED Vs DRY
F
Pyrimidine metabolism
Thymine
Uridine
beta-Alanine
Glutamine
Uracil
Purine metabolism
Glutamine
Hypoxanthine
Allantoic acid
Uric acid
Xanthine
Adenosine
Guanosine
Adenosine-5-monophosphate
Guanine
Adenine
Diguanosine tetraphosphate
DOG0 Vs DRY
DOG1 Vs DOG0
DOG3 Vs DOG1
DOG5 Vs DOG3
KILNED Vs DOG5
KILNED Vs DRY
Log2(FC)
-5
0
5
