## Supplementary Materials and Methods for "Integrative LC-MS and GC-MS Metabolic Profiling Unveils Dynamic Changes during Barley Malting"

**2.1 Plant materials, malting, and malt quality**

Equal quantities of Conrad seed grown in 2014 and 2015 at 4 barley producing regions were pooled and then malted (Table S1) at the Malt Quality Laboratory of the USDA-ARS Cereal Crops Research Unit (CCRU) in Madison, WI. Briefly, 3 micro-malting replicates of 110 g each were steeped according to the following regimen: alternating cycles of 4h water immersion at 16°C followed by 4 h air rest at 18°C repeated until the moisture content reached 45%. Post-steeping, seeds were transferred to containers with small sidewall holes and germinated for 120 h at 17°C and > 98% humidity. To prevent matting and to allow gas exchange, the containers were rotated every 30 min for 3 min and moisture levels were maintained at 45%. The germinated seeds were then kilned using the following program: 49°C for 10 h, 54°C for 4 h, 60°C for 3 h, 68°C for 2 h, and 85°C for 3 h with 30 min temperature ramps (Fig 1). After kilning the moisture content of the finished malt was 4%, the dried culms are removed, and the samples were left to rest for a week to equilibrate the moisture throughout the grains. Malt quality analysis was performed at CCRU using the CCRU methods of analysis of barley and malt (https://www.ars.usda.gov/ARSUserFiles/50900500/barleyreports/CY%20METHODS.pdf) (Table S1).

**2.2 Sample collection and grinding procedure**

9 samples were collected (three per micro-malting replicate) at 6 different stages: unmalted mature grain (DRY), out of steep/post-step (DOG0), on alternating days of germination (DOG1, DOG3, and DOG5), and at the end of kilning (KILNED) (Fig 1). Samples were collected in Eppendorf tubes, immediately flash frozen in liquid nitrogen and subsequently stored at −80°C until grinding.

Samples were lyophilized to remove moisture, enhancing the grinding efficiency and stability of the samples. 10 dried seeds were placed into 2241-PC 5 ml polycarbonate vials in a specific layering order: one 6.3 mm stainless steel ball was placed at the bottom, followed by five malted seeds, another 6.3 mm stainless steel ball, the remaining five malted seeds, and a final 6.3 mm stainless steel ball on top. Each vial was then securely capped. Prior to grinding, the vials were immersed in liquid nitrogen for 10 min to ensure thorough cooling. Subsequently, they were placed into a pre-cooled 2662 cryo-block (also pre-cooled in liquid nitrogen) to maintain low temperatures during the grinding process. The grinding was carried out using a 2010 Geno/Grinder (Spex SamplePrep) with the following program: 30 seconds of grinding at 1750 strokes per minute, followed by a 25-second rest period. This cycle was repeated three times. Immediately after the grinding program was completed, the vials were once again submerged in liquid nitrogen to rapidly cool the samples, thereby preserving their integrity until they were transferred to their final storage vials.

2.3 **Metabolite extraction**

**2.3.1 Preparation of samples for GC-MS analysis**

Finely ground samples (4 mg) were extracted using 1 ml of 3:3:2 isopropanol (IPA)/acetonitrile (ACN)/water (v/v/v) by vortexing for 10 s and shaking for 6 min at 4°C. After centrifugation at 14,000 RCF for 2 min, the supernatant was aliquoted into two equal portions and dried. One aliquot was derivatized with 10 µL of methoxamine (40 mg/mL) in pyridine and samples were agitated at 30°C for 90 minutes. Following this, 91 µL of N-methyl-N-trimethylsilyltrifluoroacetamide (MSTFA) and a mixture of fatty acid methyl esters (FAMEs) ranging from C08 to C30 was added to each sample, which was then agitated at 37°C for 30 min to complete the derivatization process. The prepared samples were then transferred to vials, sealed, and loaded into the GC-MS instrument. Quality control (QC) samples were prepared by equally pooling ground powder from all biological samples and were processed as described above. Method blanks were prepared using the same 3:3:2 IPA/ACN/water mixture without the addition of biological sample and processed using the same extraction and processing procedures as the other samples.

**2.3.2 Preparation of samples for LC-MS analysis**

Finely ground samples were extracted with 800 μl of 80% isopropanol, followed by sonication for 30 seconds, vertexing for 10 minutes and then centrifuging at 4000 rpm for 10 minutes. Subsequently, 700 μl of the supernatant was transferred to a fresh vial. This process was repeated with another 700 µl of 80% IPA, and the resulting 1.4 ml of combined supernatant was discarded. The pellet was re-extracted twice more using 700 µl of 80% methanol. The 1.4 ml of combined methanol extract was dried under nitrogen and then reconstituted in 100 µl of methanol. The reconstituted extracts were then transferred to autosampler vials for UPLC-MS analysis. QCs were prepared by equally pooling ground powder from all samples and were processed as described above. Method blanks were prepared and processed in the same manner without the addition of biological sample.

**2.4 Metabolomic Fingerprinting**

**2.4.1 GC-MS measurements**

GC-MS measurements were performed on an Agilent 7890A GC coupled with a Leco Pegasus IV time-of-flight mass spectrometer (LECO Corporation, St. Joseph, MI, USA). Derivatized samples (0.5 μl) were injected using a splitless method into a multi-baffled glass liner. Pooled QCs were injection after every 10 samples. Compounds were separated using the GC-Column DB-5MS (length: 30 m, internal diameter: 0.25 mm, film thickness: 0.25 μm, packing: 95% dime-thylpolysiloxane/5% diphenylpolysiloxane) with an Intergra-Guard at 275°C with a helium flow of 1 mL.min^−1^. The injector temperature was ramped from 50°C to 250°C by 12°C s^-1^. The temperature of the column oven was set at 50°C (held for 1 min) then ramp to 20°C/min to 330°C and then held for 5 min. The transfer line is set to 280°C while the EI source set to 250°C. The total analysis for each sample lasted 20 min. The electron ionization (EI) energy was set to 70 eV. The system collected the mass spectra in mass range of 85-500 Da at an acquisition rate of 17 spectra/s.

**2.4.2 LC-MS measurements**

One microliter of extracted sample was injected onto a Waters Acquity UPLC system (Waters Corporation, Milford, MA, USA) with a pooled quality control (QC) injection after every 5 samples. Compounds were separated using a Waters Acquity UPLC CSH Phenyl Hexyl column (1.7 μM, 1.0 x 100 mm) (Waters Corporation, Milford, MA, USA), using a gradient from solvent A (water with 0.1% ammonium formate) to solvent B (acetonitrile, 0.1% formic acid). Injections were made in 99% A, held at 99% A for 1 min, ramped to 98% B over 12 min, held at 98% B for 3 min, and then returned to starting conditions over 0.05 min and allowed to re-equilibrate for 3.95 min, with a 200 μl/min constant flow rate. The column and samples were held at 65 °C and 6 °C, respectively. The column eluent was infused into a Waters Xevo G2-XS Q-TOF-MS with an electrospray source in positive mode, scanning 50-1200 m/z at 0.1 seconds per scan, alternating between MS (6 V collision energy) and MSE mode (15-30 V ramp). Calibration was performed using sodium formate with 1 ppm mass accuracy. The capillary voltage was held at 700 V, source temperature at 150°C, and nitrogen desolvation temperature at 600°C with a flow rate of 1000 l/h.

**2.5 Data treatment and metabolite identification**

**2.5.1 GC-MS**

Following data collection, raw data files were preprocessed and stored as ChromaTOF-specific *.peg files, as generic *.txt result files, and additionally as generic ANDI MS *.cdf files. Leco ChromaTOF v2.32 was used for data preprocessing without smoothing, 3 s peak width, baseline subtraction just above the noise level, and automatic mass spectral deconvolution and peak detection at signal/noise levels of 5:1 across the entire chromatogram. Apex masses were submitted to the BinBase algorithm (Fiehn et al., 2005) using the settings: validity of chromatogram (<10 peaks with intensity >10^7 counts s^-1^), unbiased retention index marker detection (MS similarity>800, validity of intensity range for high m/z marker ions), retention index calculation by 5th order polynomial regression. Spectra are cut to 5% base peak abundance and matched to database entries from most to least abundant spectra using the following matching filters: retention index window ±2,000 units (equivalent to about ±2 s retention time), validation of unique ions and apex masses (unique ion must be included in apexing masses and present at >3% of base peak abundance), mass spectrum similarity must fit criteria dependent on peak purity and signal/noise ratios and a final isomer filter. Raw data were normalized to the sum of all peak heights for all identified metabolites, but not the unknowns, for each sample.One of the replicates from the dry seeds failed and was removed from further GC-MS analysis.

**2.5.2 LC-MS**

XCMS v3.9.3 was used for feature finding, retention time alignment, correspondence analysis, and peak filling in R v4.0.3 (Smith et al. 2006; Tautenhahn et al. 2008). XCMS steps included:

(1) peak detection using the CentWave algorithm (ppm= 30 peakwidth= c(2.2, 15), snthresh = 3, prefilter = c(3, 10), mzCenterFun = wMean, integrate = 1, mzdiff = 0.01, fitgauss = TRUE, noise = 2, verboseColumns = TRUE, roiList = list(), firstBaselineCheck = TRUE, roiScales = numeric(0), extendLengthMSW = TRUE.
(2) Peak grouping (PeakDensity) : bw = 4, minFraction = 0.5, minSamples = 1, binSize = 0.015, maxFeatures = 50.
(3) Retention time correction (PeakGroups) : minFraction = 0.5 extraPeaks = 1, smooth = loess, span = 0.2, family = gaussian, subset = integer(0), subsetAdjust = average.
(4) Peak grouping (PeakDensity) : bw = 1.75, minFraction = 0.45, minSamples = 1, binSize = 0.015, maxFeatures = 50
(5) Missing peak filling (FillChromPeaks) : expandMz = 0 expandRt = 0, ppm = 0, fixedMz = 0, fixedRt = 0.

RAMClustR v1.2.4 (Broeckling et al., 2014) in R v4.2.2 was used to normalize, filter, and cluster features into spectra. Features with missing values were replaced with small values simulating noise: for each feature, the replacement value was equal to the absolute value of 0.5 times the minimum detected value of that feature. Features were normalized by linearly regressing run order versus QC feature intensities to account for instrument signal intensity drift. Only features with a regression p-value less than 0.05 and an r-squared greater than 0.1 were corrected (32342 of 107739). Features which failed to demonstrate signal intensity of at least 2-fold greater in QC samples than in blanks were removed from the feature dataset. 25798 of 107739 features were removed. Features were further filtered based on their QC sample CV values. Only features with CV values $\leq$ 0.5 in MS or MSMSdata sets were retained (27133 of 81941 features were removed). Features were clustered using the ramclustR algorithm (Broeckling et al., 2014). Parameter settings were as follows: st = 1.99, sr = 0.5, maxt = 199, deepSplit = FALSE, hmax = 0.3, minModuleSize = 2, and cor.method = pearson. Molecular weight was inferred from in-source spectra (Broeckling et al., 2016).

MSFinder v 3.52 (Tsugawa et al., 2016) was used for spectral matching, formula inference, and tentative structure assignment. Results were imported into the RAMClustR object. A total score was calculated based on the product scores from the findmain function and the MSfinder formula and structure scores. A total of 29127 annotation hypotheses were tested for 8921 compounds. Spectra matches took precedence over computational inference-based annotations. CHEBI and COCONUT were set as priority databases, and matches to these databases were given a priority factor value of 1. Matches to databases other than the priority databases were assigned a priority factor of 0.9. A custom database of compounds found in barley was generated based on associations from FooDB and PubChem. The list of 8858 InChIKey structures in this custom barley database was also used for annotation prioritization. Annotations with InChIKey(s) that didn’t match those in the custom barley database were assigned an InChIKey priority factor of 0.9 to decrease scores for these annotations). The annotation with the highest total score was selected for each compound.

2.6 **Dataset Integration and Statistical Analysis**

We mainly focused on known metabolites (those with InChIKeys). 170 out of 773 GC-MS features and 7,600 out of 8,921 LC-MS features corresponded to known metabolites (Table S2-3). To include features with high confidence, we excluded the LC-MS features from different clusters with identical annotations. In cases, where a metabolite was detected in both datasets, we selected the data from the platform with the lower pool QC CV value (Table S4). Final integrated dataset comprised 4,980 known metabolites (Table S5). Box–Cox transformation was applied to normalize the data distribution (Yu et al., 2022).

The normalized data were then assessed for normality using the Shapiro-Wilk test. Metabolites (2,799 from the LC-MS dataset and 118 from the GC-MS dataset) that did not follow a normal distribution (p < 0.1, commonly accepted threshold for a non-normal data distribution) after the Box-Cox transformation with were subjected to the Mann-Whitney U test for comparisons between consecutive stages and the Kruskal-Wallis test for ANOVA. For metabolites with a normal distribution (2,019 from the LC-MS dataset and 44 from the GC-MS dataset), Levene’s test was used to assess the homogeneity of variances. Metabolites with unequal variances (185 from the LC-MS dataset and 9 from the GC-MS dataset, with p < 0.1) were analyzed using Welch’s t-test and Welch’s ANOVA. Metabolites with equal variances (1,833 from the LC-MS dataset and 34 from the GC-MS dataset) were subjected to Student t-test and standard ANOVA. Multiple testing using false discovery rate (FDR) correction was applied after integrating raw p-values from both LC-MS and GC-MS datasets for each statistical test analysis separately. Metabolites were considered significantly different if they met the following criteria: (i) false discovery rate < 0.05, and (ii) fold change ≥ 2 or ≤ 0.5.

Compounds were assigned to chemical ontogenies using ClassyFire (Djoumbou Feunang et al., 2016a). Chemical similarity enrichment analysis (ChemRICH) was performed using raw p-values derived from the different statistical tests applied for comparing means in consecutive stages (such as the Mann-Whitney U test, Welch's t-test, and Student's t-test) and parent class level 1 information as set definition. Sets having a frequency of less than 3 and sets having no compounds with a raw p-value < 0.10 were excluded. Sets with false discovery rate < 0.01 were considered significant. Annotated metabolite names were converted to KEGG IDs using various platforms: Refmet (Fahy & Subramaniam, 2020), Mbrole2.0 (López-Ibáñez et al., 2016), Chemical Translation Service (Wohlgemuth et al., 2010), PubChem Exchange Identifier service (<https://pubchem.ncbi.nlm.nih.gov/idexchange/>), and Metaboanalyst 6.0 (Pang et al., 2024). Out of 4,980 annotated metabolites in the dataset, 600 corresponded to unique metabolite-KEGG ID pairs. The data was autoscaled and Kyoto Encyclopedia of Genes and Genomes (KEGG) pathway analysis was performed. MetaboAnalyst 6.0 (Pang et al., 2024) was used to identify and visualize the altered metabolic pathways using *Oryza sativa* japonica as the background set. The global test and relative-betweeness centrality were used for the enrichment method and pathway topology analysis, respectively. The significantly affected pathways (with at least 5 metabolites detected in our dataset) were selected by FDR < 0.05 from pathway enrichment analysis.
